# Mechanics of *E. coli* cell width homeostasis and bulging dynamics from MreB and septum inhibition

**DOI:** 10.1101/2024.11.22.624946

**Authors:** Tanvi Kale, Ryth Dasgupta, Mandar M. Inamdar, Chaitanya A. Athale

**Affiliations:** Div. of Biology, IISER Pune, Dr. Homi Bhabha Road, Pashan, Pune 411008, India; Civil Engineering Dept., IIT Bombay, Powai, Mumbai 400076, India

## Abstract

The mechanobiology of cytoskeleton and cell envelope play a vital role in cell shape homeostasis. In the gram-negative model rod-shaped bacterium *Escherichia coli* antibiotics that weaken the cell envelope are seen to result in bulges that eventually lead to lysis. Here, we quantify the shape dynamics of *E. coli* treated with cephalexin, a PBP3 inhibitor and A22, an MreB polymerization inhibitor from single cell microscopy. We find low concentrations of both inhibitors result in bulge formation with multiple cell shapes observed: rugby, large bacilli and rods with two and three bulges. We quantify the parameters of cell envelope rigidity, pressure and fluidity obtained from fitting a computational model of shell-mechanics to length and width dynamics from experiment. Using this optimized model we estimate the turgor and growth pressures of untreated growing cells as ≈0.15 MPa and ≈0.4 MPa. The bulge expansion dynamics correlate most prominently with a change in bending rigidity of the cell wall and cytoskeleton. Simulations predict a threshold behavior in response to envelope bending rigidity that is validated by comparison with experiments of *E. coli* treated with A22 either in isolation or in combination, resulting in loss of width control and cell shape change.

## Introduction

The cytoskeleton plays both a regulatory and mechanical role in cell shape determination. In the model bacterium *Escherichia coli* both the tubulin homolog FtsZ and actin homolog MreB play vital roles in determining cell division and rod-shape, respectively. FtsZ polymers assemble to form a Z-ring, determining the site of septation (*Margolin*, 2001), while MreB assembles across the length of rod-shaped *E. coli* as multiple curved filaments essential for maintaining the rod-shape and providing sites for wall assembly (*Jones, et al.* 2001). The ‘ring like’ nature of MreB filaments formed under the inner cell membrane are thought to both detect and drive cell curvature (*Ursell, et al.* 2014). Thus MreB cytoskeletal filaments act to sense and mechanically determine the cell curvature and therefore width (*Wong, et al.* 2019). Filaments of MreB were shown to align with higher curvature regions of the cell envelope and direct cell wall synthesis (*Hussain, et al.* 2018). Thus, cell width was observed to be altered in a dose-dependent manner by MreB cytoskeletal filament filament assembly and its inhibition (*Ouzounov, et al.* 2016). MreB filament assembly has also been observed to align with the axis of greatest principal curvature in *Bacillus subtlis* with coupling to peptidoglycan synthesis that in turn reinforces the rod-shape of the cell (*Hussain, et al.* 2018). The peptidoglycan cell wall is well understood to play a vital role in cell shape maintenance with the network properties suggesting mechanisms of robustness (*Huang, et al.* 2008). The cytoskeletal and wall systems are coupled and self-organize as seen in the role of FtsZ, FtsA and ZipA assembly at the divisome site, that then drives the localization of FtsK, Q, L, W I and N (*Goehring, et al.* 2007). FtsI, a transpeptidase (Pbp3), in turn is responsible for ring constriction even in presence of FtsZ (*Pogliano, et al.* 1997). While inhibition of FtsI and related septal factors have been studied in combination, how the inhibition of these together with MreB affects cell shape and size remains less well explored.

The widely used *β*-lactam class of antibiotics work by inhibiting wall integrity. This is seen in the case of treatment with *β*-lactam antibiotics to also result in occasional central bulges in cells (*Goodell, et al.* 1976). A model of the peptidoglycan wall as an elastic network with cylindrical geometry was used to demonstrate that antibiotics that inhibited local bond formation would result in orthogonal ‘cracks’ in the shell restricted locally (*Huang, et al.* 2008). However, the cylindrical shape of the remainder of the cell would be intact and membrane and cytoplasm would expand only through the crack, resulting in a bulge. This is consistent with experimental observations of vancomycin treated cells developing bugles. Bulge formation was also observed in cells treated with an FtsI inhibitor, cephexlin, through inhibition of new peptidoglycan strands, without affecting the degradation of strands by FtsN and amidases during new wall and septum synthesis *Chung et al.* (2009). Cephalexin treated *E. coli* in timelapse microscopy demonstrated that the bulge-region primarily contains the membrane with a large variation in bulge lifetimes (*Yao, et al.* 2012). Cracks in the cell wall are indeed expected in presence of the membrane, to result in bulging due to a pressure difference (Δ*p_i_*) estimated for *E. coli* to range between 0.2 and 0.5 MPa, i.e. 2 to 5 atm using hyperosmotic solutions (*Cayley, et al.* 2000) and 3 atmosphere (0.3 MPa) using AFM measurements (*Jiang, et al.* 2011). Thus, while cell bulging of *E. coli* has been reported due to cracks in the wall, their quantitative nature, the pressure driving it and mechanical rigidity of wall and cytoskeleton required to maintain cell shape, vary across studies and methods. A comprehensive picture combining quantitative experiments and model fitting could improve our understanding of the robustness of cell shape to envelope mechanics.

The asexual growth of *E. coli* was described as a BCD cycle of birth (B), chromosome replication (C) and division (D) (*Cooper and Helmstetter*, 1968). Quantification of cell size dynamics of rod-shaped bacteria have typically measured cell length as a surrogate for size, while the width of growing *E. coli* is assumed to be constant (*Si, et al.* 2015; *Zaritsky*, 2015). The growth of *E. coli* cell length was thought to be exponential (*Cullum and Vicente*, 1978) but was more recently shown to better fit bi- or tri-linear functions (*Reshes, et al.* 2008). Bacterial cell size regulation has been modelled in terms of a ‘constant increment’ or ‘adder’ (*Amir*, 2014), consistent with cell size homeostasis (*Taheri-Araghi et al.*, 2015). At the same time, experimental tests of these models have assumed a constant width, with length extension considered to account for the increment in size (*Campos, et al.* 2014). However, since cells undergo volume growth, width increase and population variability could also confound the search for a model of bacterial cell size regulation (*Facchetti, et al.* 2019). Thus, quantification of *E. coli* width dynamics and the molecular mechanism regulating it, is vital to address the question of the principles bacterial size and shape.

*E. coli* surfacearea (SA) scales with volume (V), i.e. *SA~V^γ^*, with an exponent *γ* greater than 2/3, i.e. hypergeometric, for for multiple growth conditions (*Kale, et al.* 2023). If cell width is assume to be constant, the scaling exponent would always be *γ* ~ 0.9. However, *E. coli* is reported to have a lower scaling exponent both in our work and that from other labs (*Ojkic, et al.* 2019). Additionally, we find a growth rate dependence of the scaling exponent, which would suggest the need for a model to address not just length increment, but also the role of width. A mechanical model of cell envelope tension, bending rigidity, viscosity and pressure has predicted cell width to increase and saturate (*Banerjee, et al.* 2016). Thus, quantifying cell size dynamics and reconciling them to such a mechanical model in combination with biochemical perturbations could improve our understanding of the quantitative nature of bacterial cell shape regulation.

In this study, we used the well known *β*-lactam antibiotic cephalexin and combined it with A22, an inhibitor of MreB polymerization. While cephalexin treatment alone results in cells ‘cracking’ and A22 in rounding up of cells, the combination resulted in central, symmetric bulges with a distinct statistic arising from the rapid growth of cells. We fit the mechanical shell model to data obtained from the dynamics of cell shape parameters to quantify the mechanical properties of untreated, A22-treated and cephalexin-treated cells. We demonstrate that the cell width is not constant but increases and saturates in a manner that is primarily governed by the bending rigidity of the envelope. This in turn is strongly affected by wall and MreB mechanics, with the surface tension playing a smaller role.

## Results

### Single cell shape dynamics of *E. coli* grown in presence of MreB and FtsI inhibitors

*E. coli* cells sampled from mid-log phase cultures were grown on LB-containing agarose pads with a combination of 10 *µ*g/ml cephalexin and 2 *µ*g/ml A22, in order to affect both cell length and width respectively, with the pads embedded in a 3D printed chamber (Figure S1). The cell shapes were followed in label-free DIC timelapse microscopy for over 60 minutes. The choice of cephalexin was based on the known role of this *β*-lactam antibiotic to inhibit the transpeptidase FtsI (PBP3), preventing septal wall synthesis resulting in long filamentous cells (*Pogliano, et al.* 1997). A22 was chosen for its role in cell width regulation through MreB polymerization inhibition by monomer sequestration (*Young*, 2006) resulting in rounded up cells, described as ‘lemon’ shaped, and dose-dependent increase in cell width (*Ouzounov, et al.* 2016). MreB is a critical factor regulating cell width with mutants in the ATP-binding pocket resulting in altered cell widths (*Dye, et al.* 2011). While the combination of A22 and cephalexin has been previously reported to reduce growth rates without complete cell lysis (*Iwai, et al.* 2002), microscopic shape dynamics had not been observed. Here, we show that while cephalexin results in elongated cells (filamentation) due to a lack of septation, A22 results in ovoid (’lemon’ shaped) cells that continue to septate and divide. The combination of both results in the appearance of a central bulge, eventually leading to lysis after 120 min (Figure 1A). Filamentation results from inhibition of peptidoglycan crosslinks by cephalexin, while the roundingup of cells is due to inhibition of MreB assembly. The central bulge is thought to be the combinatorial result of mechanical weakening of the wall in the central region to due to continued activity of peptidoglycan hydrolases, the absence of crosslink synthesis and the absence of MreB polymers, required to reinforce the cell wall and maintain cell width (Figure 1B).

**Figure 1:**
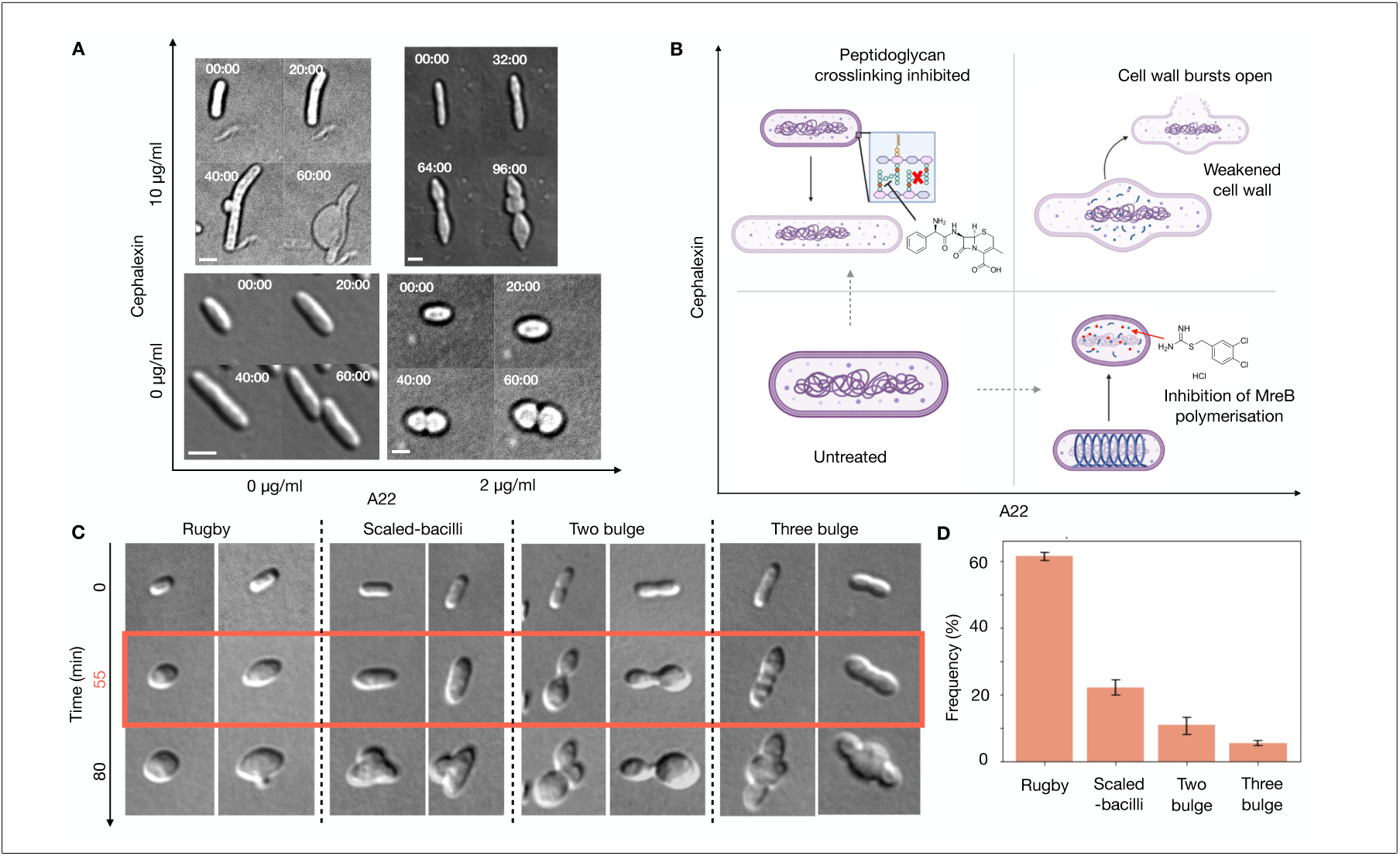
Diversity of *E. coli* cell shapes treated with A22 and cephalexin. (A,B) The effect of A22 and cephalexin on cell shape are represented in terms of their individual and combined effects from: (A) DIC microscopy of *E. coli* MG1655 comparing the shapes of cells that were *(lower left)* untreated, or treated with either *(upper left)* 10 *µ*g/ml cephalexin, *(lower left)* 2 *µ*g/ml A22 or *(upper right)* a combination of both. Scalebar: 1*µ*m. (B) The schematic represents the potential mode of action with *(upper left)* cephalexin alone inhibiting cell septum synthesis and cell filamentation, *(lower right)* A22 alone inhibiting of MreB polymerization and resulting in ellipsoid cells and *(upper right)* the combined treatment results in in bulged cells due to weakening of the cell wall at the center along with lack of mechanical support due to inhibition of MreB polymerization. (C) Representative images of cells treated with 10 *µ*g/ml cephalexin + 2 *µ*g/ml A22 at time points 0, 55 and 80 min of treatment are classified into distinct morphologies as: Rugby, Scaled-bacilli, Two bulge and Three bulge. (D) The frequency of cell-shapes is plotted as a percentage of the total. Error bar: s.d. Biological replicates n=3, with the number of ROIs analyzed 10 (*n_cells_*= 347), 4 (*n_cells_*= 356) and 10 (*n_cells_*= 298).

We find that in addition to these representative bulges, cells take on varied shapes that also, in turn, changing in time over ~90 minutes after growth on agarose-pads containing the inhibitors. We classify these shapes as: (i) rugby, (ii) scaled-bacilli, (iii) two-bulge, (iv) three-bulge (Figure 1C). Rugby-shaped cells are similar to what has been described as ‘lemon’ shaped, while ‘scaled bacilli’ refers to a proportionate increase in both length and width. The presence of multiple bulges appears qualitatively to correspond to exposure of cells just before the onset of cell division. We find ‘rugby’ and ‘scaled-bacilli’ are most frequently observed in our experiments, as compared to the 2- and 3-bulge shapes with over three biological replicates, with 347, 356 and 298 cells analyzed in each (Figure 1D). We believe the heterogeneity in shapes arises from the unsynchronized nature of the sampling method. The statistics demonstrate that the rugby shaped cells are most frequent. This can be understood in the sense that mid-log phase cultures sampled have a majority of newborn cells, i.e. cells in the birth stage. Such cells when exposed to low concentrations of both cephalexin and A22, continue to grow in volume without the ability to synthesise new cell wall and are unable to maintain cell width due to the inhibition MreB polymerization. Since at this stage, the septum formation has not been initiated, the cell wall in the central region of the cell remains unaffected. Cells that have been categorized in to the ‘two bulge’ category, are likely to be the result of mid-cell envelope rigidity being maintained in the late division stage. However, as the volume grows, without new cell wall synthesis from the new two bulges in the regions which are weaker maybe observed. This is confirmed by observing the slight constriction at the mid-cell region at t = 0, indicating septum formation. Some cells appear to form ‘scaled bacilli’, i.e. increase in width due to MreB assembly inhibition, while cell elongation continues. Some cells are observed to lyse from the mid-cell region if the the FtsZ ring positioning has happened and septum formation initiated. In cases where the daughter cells were about to form, i.e. division (D) stage, but the septum was inhibited, while cell elongation continued, with the next round of chromosome replication (C) being entered, ending with yet another septum being inhibited-thus giving rise to initially two bulges and finally weakening of the initial septum - giving rise to 3 bulges. The bulging itself appears to be the result of the combined action of amidases which break the peptide bonds, without the transpeptidase to rebuild the gap, and the absence of MreB to maintain curvature of the cell envelope.

We also demonstrate that this treatment of cephalexin and A22 does not affect membrane integrity, as observed in the continuous label of the membrane in cephalexin-(Figure S2A), A22-(Figure S2B) and A22+cephalexin - treated cells (Figure S2C). Thus, the change in cell shape is likely to be mechanical, and not due to membrane disruption.

Since MreB and cell wall integrity are known to play a vital role in cell mechanics, we proceeded to model the mechanics of the cell envelope and compare them to experiment.

### Optimizing mathematical model of single-cell length-width growth dynamics to estimate cell mechanics

The mechanics of a growing rod-shaped cell like *E. coli* has been previously modeled as a shell with the equation of motion used to formulate two ordinary differential equations (ODEs) for the time-evolution of length (L) and width (w) in the small deformation regime (*Banerjee, et al.* 2016). These ODEs relate the internal energy for a rod-shaped bacterium (*E_rod_*) to the dissipative forces associated with degrees of freedom, length and width, resulting in the dynamics of length being described by the equation:

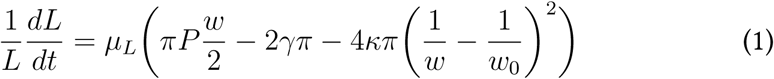

and of cell width by:

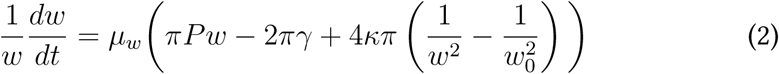

Here, *P* is the total pressure combining the effects of turgor and growth, *w* is the width of the cell, *L* is the length of the cell, *γ* the surface tension of the cellular envelope, *κ* the radial bending rigidity of the combination of MreB, cell wall and membrane, *w*_0_ the preferred width and *µ_L_* and *µ_w_* the mobility of the shell along the length and width respectively (Figure 2A, B). The mobility is inversely related to the effective viscosity for length as *µ_L_* = 1*/*2*πhη_L_* and along the width by *µ_w_* = 1*/*2*πhη_w_*. Here, *η_L_* and *η_w_* are the viscosity along length and width respectively to account for anisotropy of envelope extension along the length and width respectively. Here, *h* = 3 nm is the thickness of the envelope.

**Figure 2:**
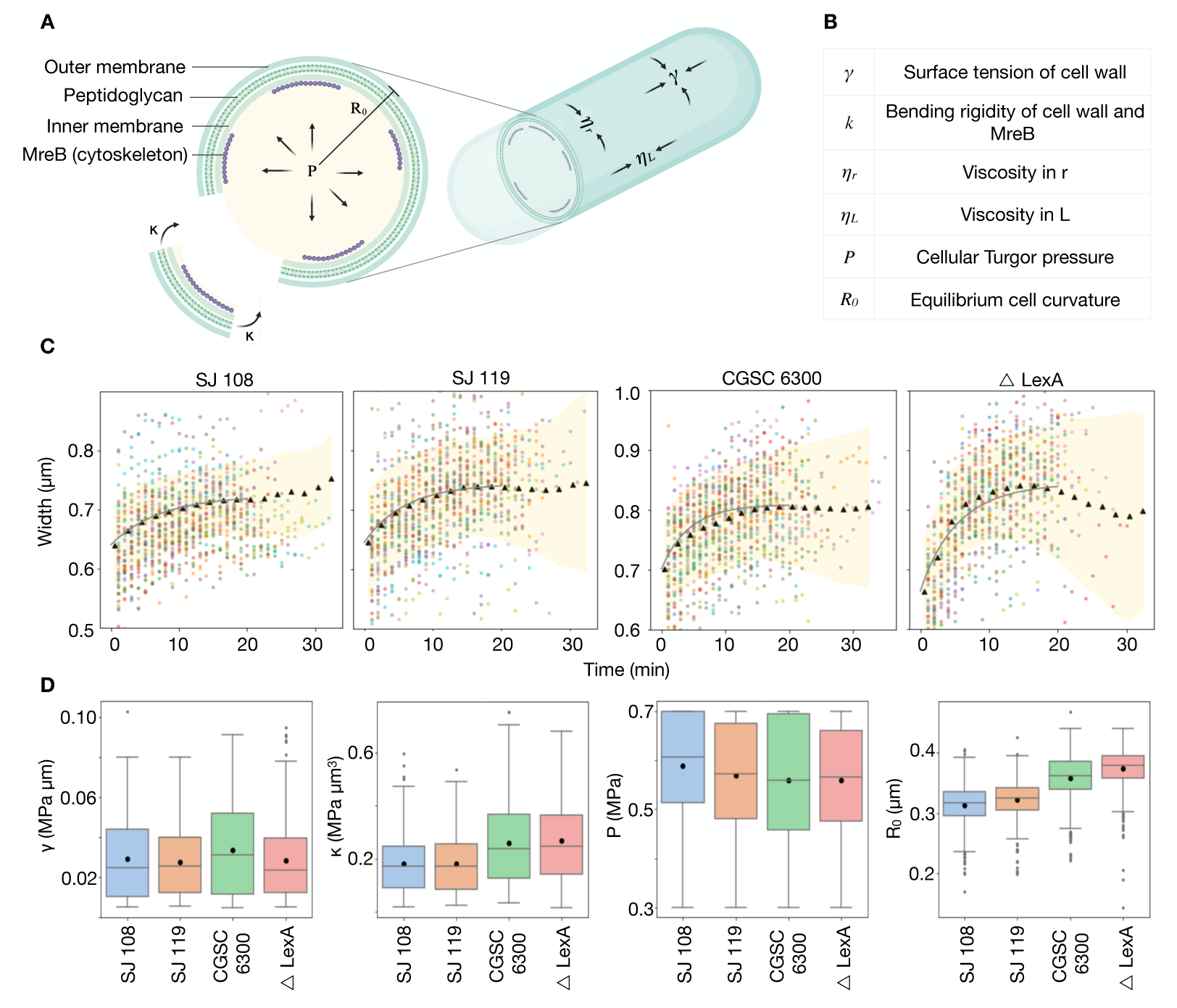
Fitting a model of cell mechanics to experimental data of *E. coli* growth dynamics. (A,B) The shell-model of cell growth based on the mechanical properties of cell wall surface tension, bending rigidity, viscosity and the turgor pressure of the cell described previously (*Banerjee, et al.* 2016) is (A) schematically represented with (B) the parameters described in a table. (C) The two-ODE model simulations were fit to the dynamics of cell widths and lengths (Supplementary Figure S4) obtained from previous experiments (*Wang, et al.* 2010) with colored circles: individual cell width dynamics, black triangles: binned averages for a constant bin size (2 min), shaded yellow region: s.d. of the mean and black line: fit to the average data. (D) The estimated parameters from model fitting are compared across strains (colors) in terms of surface tension, *γ* (MPa *µ*m); bending rigidity, *κ* (MPa *µm*^3^); total pressure, P (MPa) and equilibrium cell curvature, *R*_0_ (*µ*m). Black circle: mean, horizontal line: median, box: first and third quartile, gray dots: outliers.

The ODEs were numerically solved and the model parameters optimized by fit to experimental measurements of length and width dynamics using particle swarm optimization (PSO), as described in the Materials and Methods section. In order to estimate the mechanical fit-parameters of untreated cells, a previously described dataset of multiple ‘wild type’ *E. coli* strains grown in a microfluidics ‘mother machine’ (*Wang, et al.* 2010) was parsed, pre-processed and pruned to remove spurious values of lengths and widths (Figure S3). Such sub-sampled data was averaged and the simulations fit simultaneously to cell width (Figure 2C) and length dynamics (Figure S4). The length-width dynamics of n = 500 cells randomly sub-sampled from a larger dataset of 10^4^ cells were individually fit to obtain statistics in the fitting parameters (Figure 2D). While some of the fitting parameters are different between strains, the differences do not alter cell sizes and shapes, suggesting a degree of robustness to variations in the mechanical parameters. For ‘wild type’ cells the average surface tension *γ* ∼ 0.03 MPa *µ*m, bending rigidity *κ* ∼ 0.22 MPa *µ*m^3^, total pressure *P* ∼ 0.56 MPa and equilibrium width *w*_0_ ∼ 0.7 *µ*m. The pressure and bending rigidity are comparable to those reported in literature (Table S1). Thus, we demonstrate quantification of cell mechanics by model optimization to microscopy data. While individual model fits are robust, variability in fit parameters for each strain measured by the coefficient of variation (Figure S5) could be the result of heterogeneity in individual cell widths (Figure S6).

We proceed to examine if this model can be used to both expand our understanding of the mechanics of growing bacteria and quantify the thresholds of mechanics that alter cell shape.

### Estimating turgor pressure by minimizing volume growth in the model

The total cellular pressure obtained from our model-based fitting compared to literature revealed that most quantify turgor pressure (*P_t_*). The turgor pressure using hyperosmotic solutions was found to be 0.2 to 0.5 MPa (*Cayley, et al.* 2000) and 0.3 MPa using AFM (*Jiang, et al.* 2011). In order to make an exact comparison to our model-fitting estimates, we needed to estimate the turgor pressure. To this end, we simulated the model described in Equations 1 and 2 for P ranging from 0.01 to 0.7 MPa to find the pressure at which the cell neither grows nor shrinks. In the simple shell model this is equivalent to growth pressure *P_g_* = 0. Since the total pressure is the sum of turgor and growth pressures, i.e. *P* = *P_g_* + *P_t_*, this pressure of no growth would then be the turgor pressure *P_t_*. Growth was simulated resulting in cell width (Figure 3A) and length dynamics (Figure 3B), which in turn were used to calculate cell volume dynamics (Figure 3C) based on the spherocylinder approximation of volume, with *V* = *πr*^2^*L* + <u>^4^</u> *πr*^3^, where *L* is the length of the cell and *r* is radius which is *w*/2 where *w* is the width. Cell growth is defined as volumetric change *dV/dt* from the discrete derivative of the time-dependent change in volume. Thus when dV/dt=0, the cell neither grows nor shrinks (Figure 3D). The maximal change in volume growth max[*dV/dt*] (units *µ*m^3^/min) is used as a characteristic value for whether the cell volume shrinks, grows or undergoes no-change. Indeed, we observe a value of pressure where max[*dV/dt*] = 0 *µ*m^3^/min when input pressure is 0.15 MPa (Figure 3D). Based on the conservation relation, in the absence of growth pressure this is the turgor pressure *P_t_* of 0.15 MPa or 1.5 atm. Taken together with our fit-based estimate of the total pressure (*P*) of 0.56 MPa or 5.6 atm for growing cells (Table S1), the mean growth pressure (*P_g_*) is 0.41 MPa or 4.1 atm (Figure 3E).

**Figure 3:**
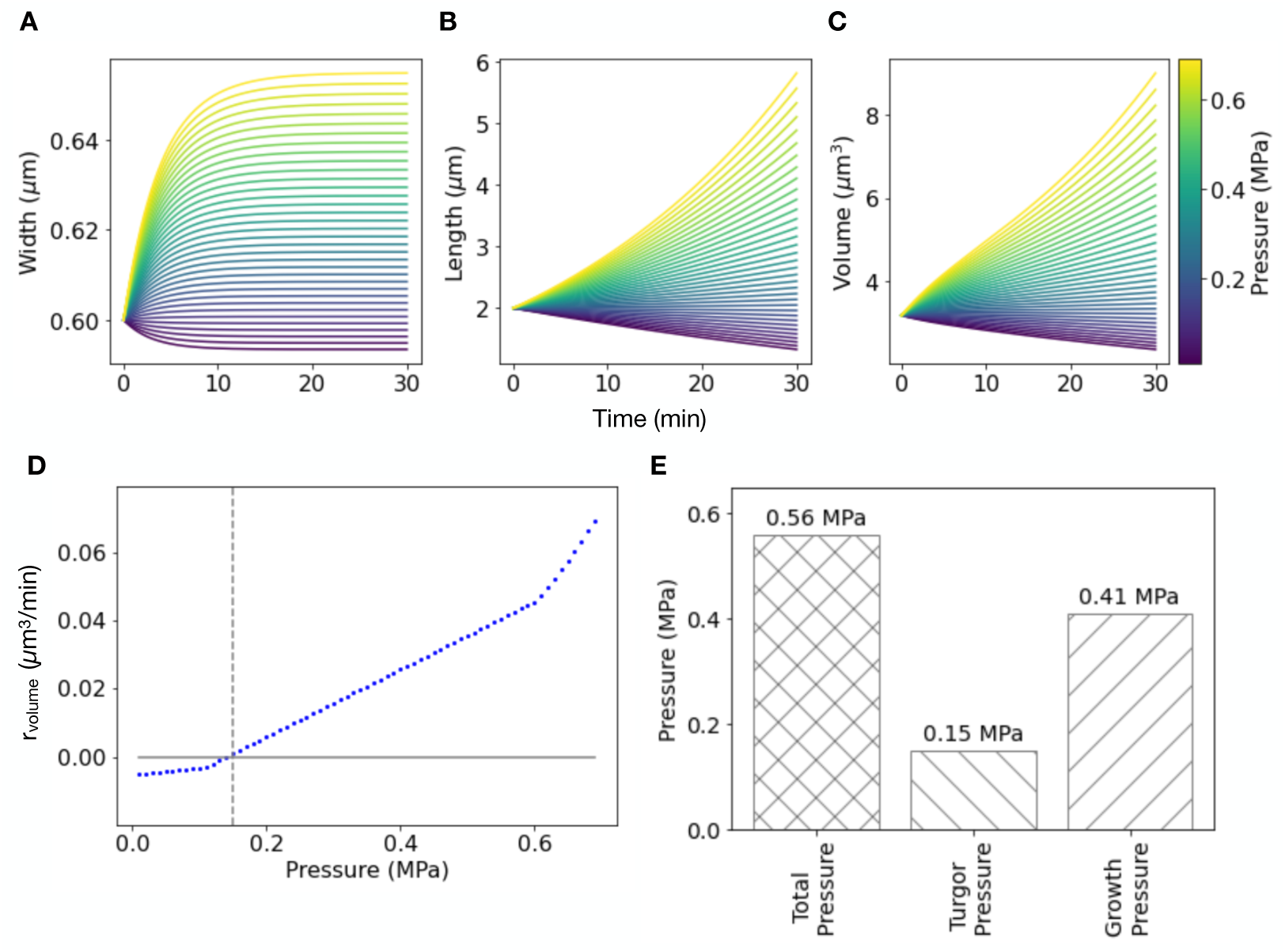
Estimating the turgor pressure and growth pressure from simulations. (A-C) The coupled ODE model (Equations 1, 2) was solved for increasing pressure, *P* (color bar). The resulting dynamics of (A) length, (B) width and (C) calculated volume with time are plotted. All other parameters were constant, inferred from the averages for all the *E. coli* strains fit to the model in Figure 2C,D (Table S1). (D) The maximal rate of cell volume growth, max[*dV/dt*] (blue circles) from simulations is plotted as a function of increasing pressure (x-axis). Solid gray line: pressure at which max[*dV/dt*] = 0. (E) The estimated total pressure and the turgor pressure (pressure at which dV/dt=0) are used to calculate the growth pressure.

With these results in hand of the turgor and growth pressure, we proceeded to apply this model-based fitting approach to quantify the mechanical effect of cephalexin and A22 in combination, and their correlation with cell shape.

### Effect of A22 and cephalexin treatment on *E. coli* cell envelope surface tension and bending rigidity

We proceeded to apply the simple rod-shaped cell model to estimate the mechanical parameter values associated with cell bulging on treatment with cephalexin and A22 (Figure 1A). Additionally since growth itself does not seem to be hindered by A22 or cephalexin alone, we also assumed a. The growth dynamics of cells cotreated with A22+ cephalexin demonstrate a distinct pre-bulge and post-bulge rate with an inflection, corresponding to the formation of a bulge (Figure S7), and so the pre- and post-bulge dynamics optimized by two distinct model parameter sets. In initial optimization of the ODE model with free pressure and equilibrium width (*w*_0_) we find A22+cephalexin treated cells have a slightly lower pressure compared to untreated due to the increasing larger cell wall area (Figure 4C, pre-bulge, post-bulge). Comparison of data across single cell time series is based on virtual alignment by the bulging inflection point, resulting in two phases which correspond to different mechanical parameters from the fits: pre-bulge and post-bulge (Figure S8). The viscosity terms were varied for post-bulge conditions, to obtain a good fit to the width data. The *w*_0_, the preferred width, is comparable for all conditions except for the A22 and cephalexin co-treatment (Figure 4D). Since the material properties of the wall and membrane are not altered the viscosity (*η_L_* and *η_w_*) was kept constant, while total pressure *P* was assumed to relate to cell growth status and environmental osmolarity and also assumed to be constant, thus reducing the free parameters to the surface tension (*γ*) and bending rigidity (*κ*). We find the surface tension to be significantly higher in +A22 treated cells compared to untreated while all under conditions are similar in value (Figure 4E). This may be attributed to a combination of reduced cell wall synthesis in absence of MreB polymers, while cytoplasmic volume increases, resulting in heightened tension on the envelope. As expected +A22 cells have a reduced bending rigidity as compared to control based on the canonical role of MreB in cell width and curvature regulation, also seen in +A22+cephalexin and pre-bulge cells (Figure 4E). This diminished bending rigidity implies that the cell width is easily deformed. The higher bending rigidity in post-bugle cells is an artifact of the rod-shaped cell model which assumes the whole cell width has expanded.

**Figure 4:**
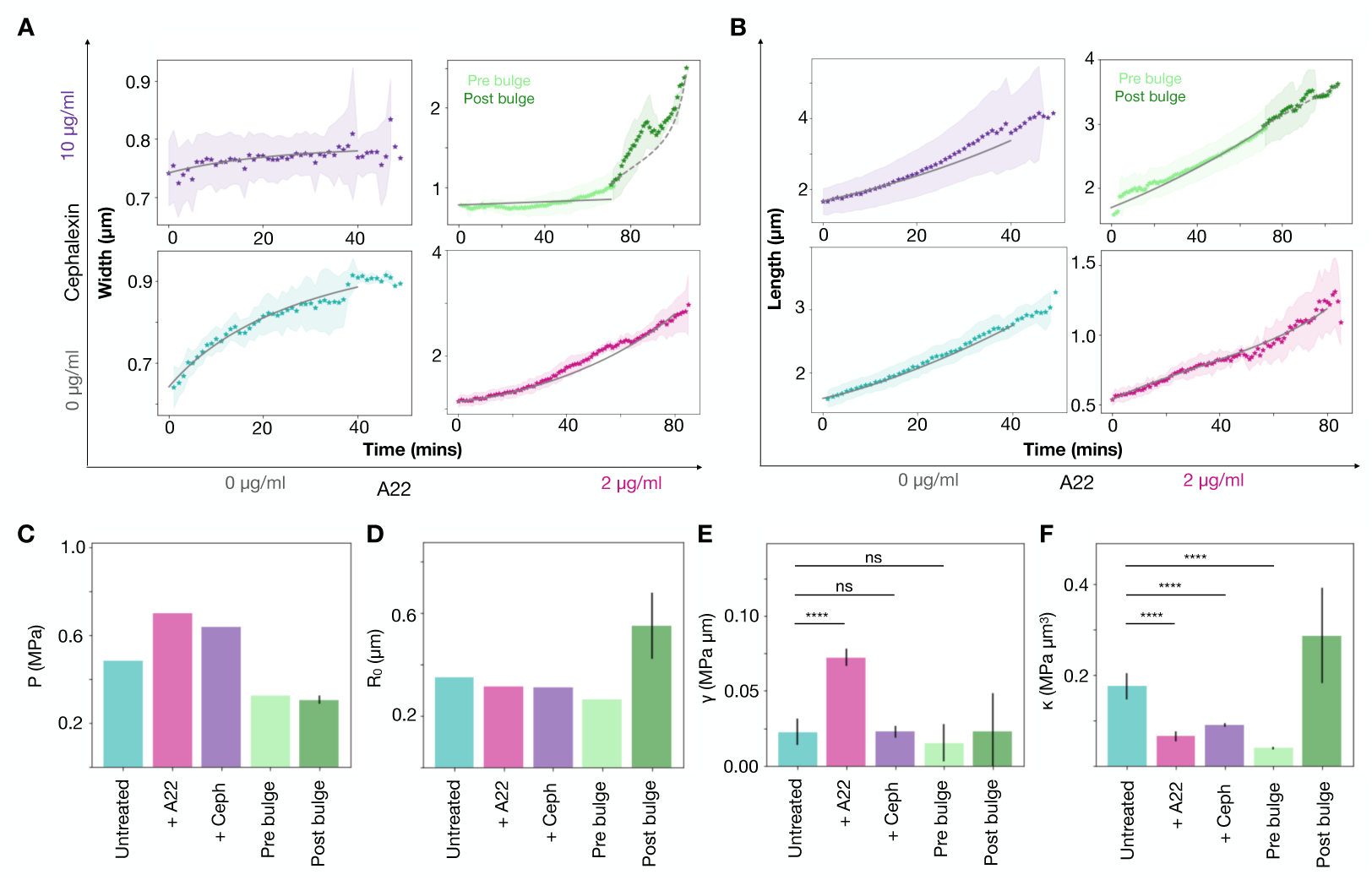
Altered bending rigidity and surface tension inferred from drug-based cell shape perturbation. (A) Width and (B) length dynamics of cells (circles: mean, shaded area: s.d.) of *E. coli* MG1655 cells were fit to the ODE model (gray line). Cells were either *(lower left)* untreated or treated with *(top left)* 10 *µ*g/ml cephalexin, *(lower right)* 2 *µ*g/ml A22 or *(top right)* a combination of both. Dashed lines: fit to the post-bulge time-points. (C-F) Model based fit parameters are compared across treatments in terms of (C) total pressure, P (MPa), (D) equilibrium cell curvature, *R*_0_ (*µ*m), (E) surface tension, *γ* (MPa *µ*m) and (F) bending rigidity, *κ* (MPa *µm*^3^). Colors correspond to conditions with cyan: Untreated, magenta: +A22, violet: +ceph, light green: pre bulge and dark green: post bulge both treated with +A22+ceph. Error bars: s.d., n*_untreated_* = 8, n_+_*_A_*_22_ = 6, n_+_*_ceph_* = 10, n*_prebulge_* = 10, n*_postbulge_* = 10. Asterisks indicate * p *<* 0.05, ** p *<* 0.01, *** p *<* 0.001, and **** p *<* 0.0001, while ns = not significant for Welch’s t-test comparing treatment pairwise with untreated cells.

We conclude that the mechanical shell model predicts that the width of cells that maintain their rod-shape geometry follows saturation kinetics. We then proceed to further use a combination of the energetics model along with a simple phenomenological model to explore the range of mechanical parameters within which the cell width is robustly maintained.

### Testing phenomenological models of *E. coli* cell width dynamics in single-cell growth

The role of a cell-size regulation involves how exactly the length and width of *E. coli* cells may be coupled to change with time. The length growth of rod-shaped bacteria has been reported variously to be either exponential (*Cullum and Vicente*, 1978; *Wang, et al.* 2010) or bi- and tri-linear (*Reshes, et al.* 2008). To address this, we formulate three alternative phenomenological models of *E. coli* width regulation: I: constant cell width resulting from intrinsic limits in width regulation which can be modeled as:

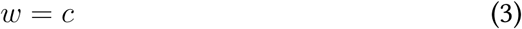

where *c* is a constant determined potentially set by the interplay of membrane precursors and MreB polymers. II: constant aspect ratio due to a proportionate increase in cell width correlating with cell length described by:

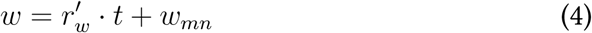

where 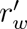 is the rate of width growth, *t* is the time in minutes and *w_mn_* is the minimum width. Such a model assumes a linear increase in cell length and ideal geometric growth (*Kale, et al.* 2023). III: width saturation which shows an initial rapid increase followed by a steady state, given by:

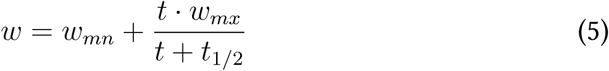

where *w_mx_* and *w_mn_* are the normalized maximal value and the minimum value of the cell width, *t* is the time during growth and *t*_1_*_/_*_2_ is the time at which the width is half maximal. The model assumes that cell width increases due to the elasticity in the peptidoglycan network and bending of MreB filaments (Figure 5A, B).

**Figure 5:**
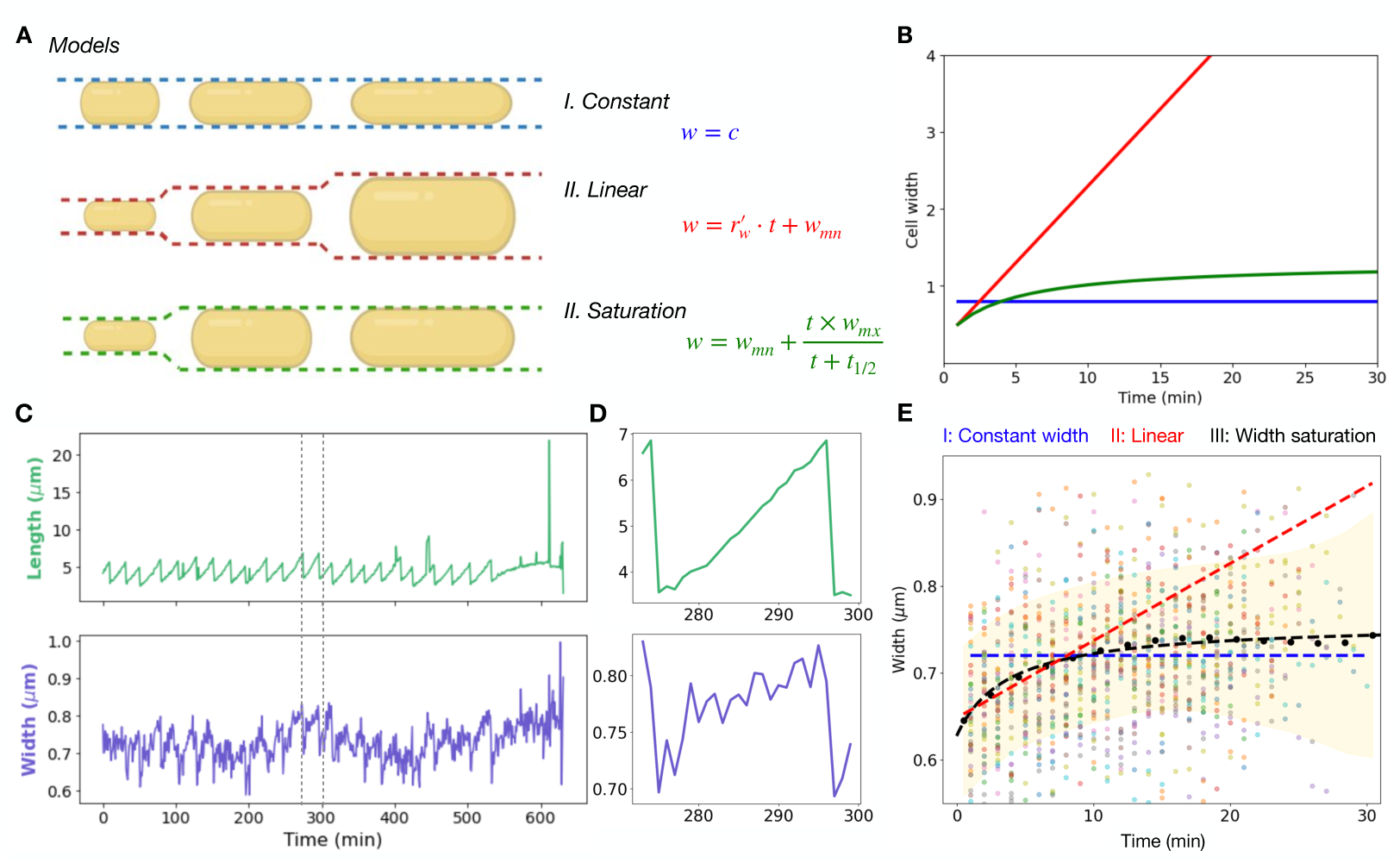
Comparing alternative phenomenological models of width growth to experiment. (A) Cell width (*w*) change with time (*t*) is modeled phenomenologically as either I: constant (blue), II: linear (red) or III: saturation (green). *c*: constant width, *r_w_^′^*: rate of width increase, *w_mn_*, *w_mx_*: minimal and maximal widths. Cells are depicted in yellow, dashed lines represent width change. (B) Representative width dynamics with time predicted by the models are graphed. Model parameters used are: (I) w = 0.8 *µ*m, (II) *A_R_* = 5 (*r_w_^′^* = 1/*A_R_* = 0.2), *w_mn_* = 0.3 *µ*m and (III) *w_mn_* = 0.3 *µ*m, *w_mx_* = 1 *µ*m and *t*_1_*_/_*_2_ = 4 min. (C,D) Experimental growth dynamics of single *E. coli* SJ119 cells from published work (*Wang, et al.* 2010) are plotted in terms of cell lengths and width as a function of time for (C) multiple rounds of division and (D) a single cell division. (E) This data is pooled from multiple cells (n = 500) and their widths plotted with time (colored circles) and averaged (black circles). The models were fit to the average data by either I: constant width (Equation 3) with c = 0.7 *µ*m (*R*^2^ = −0.62, blue), II: linear width increase (Equation 4) with *r_w_^′^* = 0.09 *µ*m/s and *w_mn_* = 0.65 *µ*m (*R*^2^ = −10.05, red) or III: width saturation models (Equation 5), with *w_max_* = 0.7 *µ*m, *w_min_* = 0.68 and *t*_1_*_/_*_2_ = 3 min (*R*^2^ = 0.97, black).

The lengths and width dynamics of *E. coli* single cells in the mother-machine from a previous report (*Wang, et al.* 2010) demonstrate successive cycles of elongation and width increase (Figure 5C). Individual dynamics of any one cell suggest that cell length increases linearly, while the width dynamics of individual cells appear more ‘noisy’, possibly due to spatial resolution of the measurement (Figure 5D). In order to compare the phenomenological models of width growth, we average the width of n = 500 cells from the previous measurements (Figure 2C) and find the data of all four strains is best fit by the width saturation model, Model III (Figure 5E). While the constant cell width (model I) model fits poorly (*R*^2^ = −0.62 for constant w = 0.7 *µ*m), the linear width growth (model II) yields a *w_min_* = 0.65 and a very high aspect ratio ( 10^2^) based on fitting the data from 0-10 minutes in which the width increases. The linear model assumes a constant aspect ratio (L/w), which would result from geometric growth which is constant and linear for length and width. However, this assumption is physiologically unrealistic, since this would imply that both width and length grow at the same rate. Thus both models I and II are rejected based on comparison to data as well as first principles.

The saturation model (model III) has three parameters: *w_mx_*, *t*_1_*_/_*_2_ and *w_mn_* (model III). Their distributions from n ~ 1000 show similar mean half-maximal time (*t*_1_*_/_*_2_) between strains (Figure S6B), while the maximal width (*w_mx_*) does appear to show small strain-dependent differences (Figure S6C). The similarity in *t*_1_*_/_*_2_ suggests that the process driving width growth has similar kinetics at a mechanistic level-total cellular pressure, cell wall growth and MreB assembly dynamics.

Therefore, we conclude that of the models tested, the width saturation model (model III) best explains the experimental cell width growth dynamics. We proceed to integrate this model with the ODE model to quantify the effect of mechanical properties on cell width growth.

### Comparing the effect of bending rigidity and surface tension on cell shape and width regulation between simulation and experiments

In order to infer possible mechanical limits to robust cell width regulation, we combined the phenomenological model with the coupled ODE model and proceeded to compare these outputs with the experiments.

We systematically *simulated dynamics of width growth* resulting from the coupled ODE model (Equations 1, 2) for varying, biologically relevant range of bending rigidity (*κ*) and surface tension (*γ*) (Figure 6A). In order to capture the kinetics of the simulated width dynamics we aimed to use a phenomenological model of width increase, similar to that described in the previous section. However, cells treated with A22 either by itself or in combination with cephalexin result in width dynamics that does not saturate (Figure 4(A)), and thus cannot be fit as well as untreated, by the width saturation model (Equation 5). Therefore, we chose to fit to a more general model of logistic growth (Equation 8). We confirm that this model choice does not alter the *w_mx_* values obtained by fitting untreated cells with the saturation model resulting in *w_mx_* of 0.75 *µ*m and the logistic model resulting in *w_mx_* of 0.74 *µ*m (Figure S9, Table S3). We therefore parametrized the *w_mx_* from simulations for a range of *κ* of 0.01 to 0.2 MPa *µ*m^3^ and *γ* ranging between 0.01 and 0.08 MPa *µ*m and fit the values to a Poisson distribution (Figure 6B, *inset*). Only those width dynamics fits from simulations that had *R*^2^ *>* 0.8 in the logistic function fit were considered in the distribution. We then divided the phase-space defined by *κ* and *γ* into three regions based on thresholds that correspond to *w_mx_* values: the mean 0.83 *µ*m (white dashed line) and maximum 1.3 *µ*m (gray dashed line). We hypothesize that cell widths are regulated in Region 1 (*w_mx_* ≤ 0.83 *µ*m), while in Region 2 (1.3 *w_mx_* 0.83 *µ*m) the width increases in an unregulated manner. The cell width cannot be determined by our model in Region 3 (1.3 *> w_mx_*).

**Figure 6:**
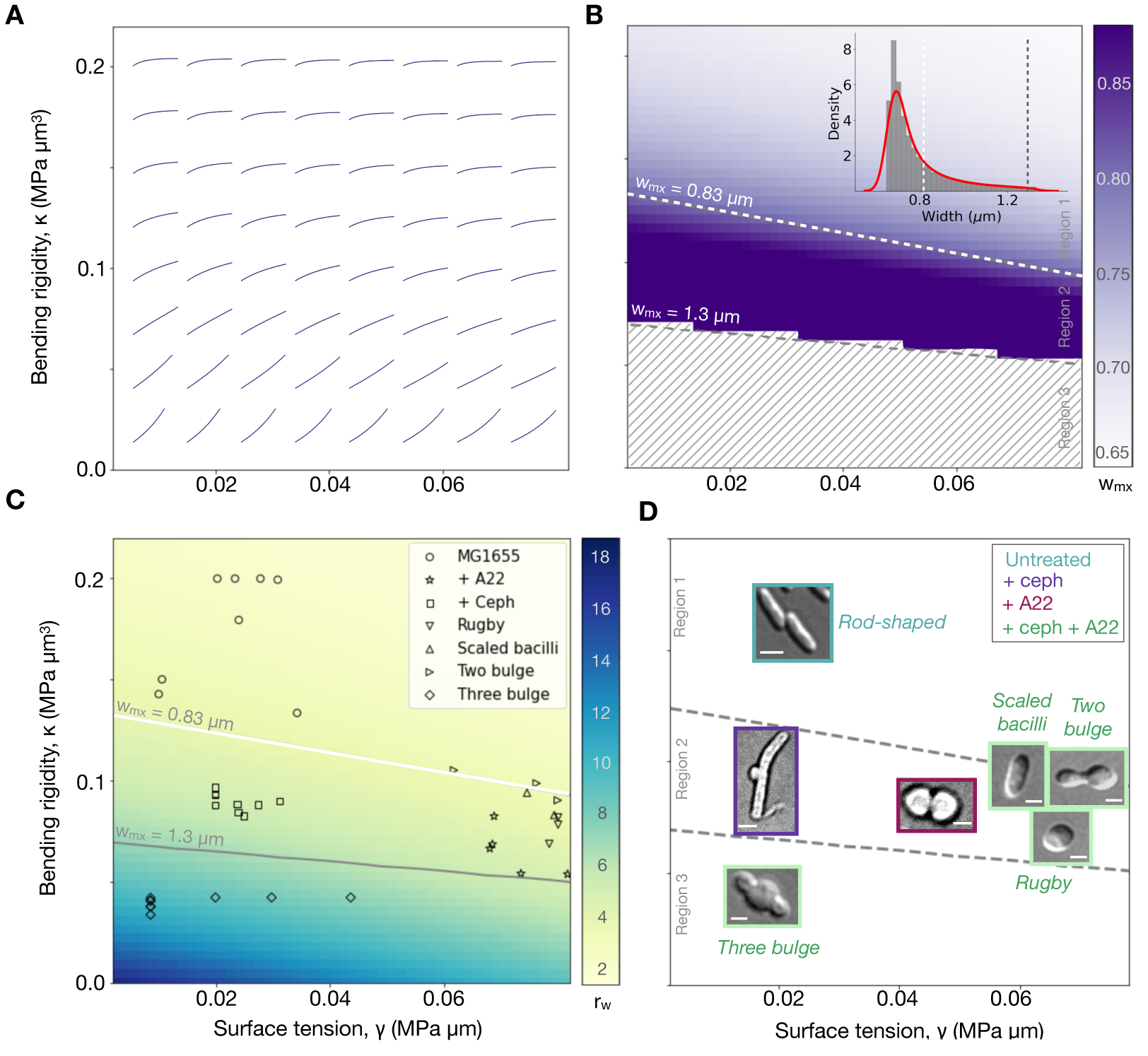
Comparing width dynamics in simulations to experiments. (A) Simulation outputs of the ODE model of shell mechanics predicts the dynamics of width growth (lines) with time (0 to 30 min) for a range of values of envelope flexural rigidity (*κ*) and surface tension (*γ*). (B) The maximal width, *w_mx_*, obtained from fitting the width saturation model (III) to the dynamics in (A) is observed in the 2D plot, with the mean of all *w_mx_* (dashed white line) used to separate it into Region 1 and 2. The hatched space represents poor fits to the model, when the R^2^ *<*0.8 and the bounding value of *w_mx_* = 1.3 *µ*m (dashed gray line). *(Inset)* The distribution of all *w_mx_* values with the mean (white dashed line) *w_mx_* = 0.83 *µ*m and upper limit to *w_mx_* (gray dashed line) *w_mx_* = 1.3 *µ*m. Colorbar: *w_mx_* (*µ*m). (C, D) The experimental fit values of *κ* and *γ* obtained for *E. coli* MG1655 cells treated with cephalexin and/or A22 are used to plot them in the same phase-space with the boundaries set by the *w_mx_* thresholds from (B) with (C) symbols representing individual cells and their shapes and (D) representative images of the cells.

Along with the changing maximum width, we predict that the rate at which the width grows also changes through the phase space. In order to calculate this rate, we used numerically differentiated the cell width time series to calculate the average rate of width growth:

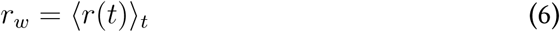

and graphed as a function of the bending rigidity and surface tension (Figure 6C). The width growth rate *r_w_* is a representative value for how fast the width is changing for untreated cells as compared to cells treated with inhibitors, which lack width regulation with values ranging from ≈2 nm/minute to ≈ 18 nm/minute. We observe that the width growth rate in **Region 1** is uniformly low (~2 nm/min) indicating that the width is well regulated. In **Region 2**, we observe that the rate of width growth is slightly higher (~4 to 7 nm/min) and changes through the phase-space. The region where the *r_w_* changes dramatically is **Region 3** (~7 to 18 nm/min) where all the factors governing width control and deregulated.

When the experimental data was overlaid on this phase space using the *κ* and *γ* values obtained from the model fits, we observe that the data for the untreated cells whose lengths are not perturbed fall into **Region 1**. The +A22 or +cephalexin data lies in **Region 2** (Figure 6C) while co-treatment data lies in **Region 2** or **3**, depending on the integrity of the cell wall. With very low *γ* and *κ* values, the three bulge data falls in **Region 3** since both the surface tension and bending rigidity are compromised due to the rupture of the cell wall. The cells with the rugby, scaled bacilli and the two bulge morphologies lie in **Region 2**. Hence, the distinct cell morphologies which are a result of the different perturbations are discretely separate in different parts of the *κ*-*γ* phase space (Figure 6D).

A further prediction of the model is that such rod-shaped cell growth will be well regulated for a threshold relation between bending rigidity and surface tension as follows:

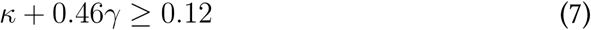

Comparing the model predictions to experiments with *E. coli* suggests that the thresholds of *κ* and *γ* could be predictive of more general mechanical limits to cell width regulation in rod-shaped bacteria due to the critical role of MreB upstream in the maintenance of envelope integrity during growth.

## Discussion

Using a combination of quantitative DIC microscopy of *E. coli* cells and a mechanical model of envelope rigidity, we have estimated the turgor and growth pressures of untreated cells along with bending rigidity and surface tension of the combined cell wall and cytoskeletal system that appears to govern cell shape. We show cells treated with both cephalexin and A22 undergo bulging and show multiple shapes that reflect their growth stage. The mechanical properties of these aberrant cell shapes suggest that bending rigidity appears to be the most critical to maintaining cell shape. The experimentally observed heterogeneity in cell shape correlates with changes in both bending rigidity and surface tension of the cell envelope. We find the predicted boundaries of cell shape maintenance in terms of bending rigidity and surface tension are validated by experimentally observed deviations from rod shapes.

Our observations of cell filamentation by cephalexin are consistent with its role as an inhibitor of septum assembly through the inhibition of FtsI (*Pogliano, et al.* 1997), while the ellipsoidal cell morphology with A22 treatment is also consistent with the inhibition of MreB resulting in a loss of rod-shape (*Ouzounov, et al.* 2016). A combination of both A22 and cephalexin was reported previously not to lyse cells (*Iwai, et al.* 2002) but the effect of the combination of both inhibitors in cell morphology had not been reported before.The diversity in cell shapes that we observe as a result of this co-treatment, could we believe be of the different cell cycle stages at the time of treatment. The quantification of the frequency of the different shapes shows that rugby-shaped cells are the most abundant. We hypothesize that these are the result of exposure of our sampling of a mid-log phase cultures, in which newborn cells are most frequently observed due to the rapid growth. As a result the effect of A22 dominates, inhibiting MreB polymerization and resulting in the observed shape due to increased flexibility in the combined envelope, i.e. membrane, wall and cytoskeleton. We could in future test this by treating synchronized cells or using pulsed flows in microfluidics.

To further investigate the mechanics behind *E. coli* cell size regulation, we used a previously described energetics and dissipation based model (*Banerjee, et al.* 2016) incorporating factors such as turgor pressure, surface tension, and bending rigidity. Previous studies have reported turgor pressures ranging from 0.03 MPa (measured via atomic force microscopy) to values as high as 0.5 MPa based on osmotic pressure studies in different growth conditions (*Deng, et al.* 2011; *Cayley, et al.* 2000). Our model fitting based estimate of turgor pressure of 0.15 MPa is within this range, and non-invasive based only on fitting the kinetics of growth. Notably, our surface tension estimate of *γ* = 0.032 MPa *µ*m is in agreement with a previously reported value of 0.019 MPa *µ*m (*Deng, et al.* 2011). In future we expect that such a fitting approach could be used to further screen for the quantitative effect of mutations or inhibitors on cell envelope mechanics. Building on these insights, we applied an energy minimization approach, similar to that of Banerjee et al. (*Banerjee, et al.* 2016), to quantify how bending rigidity and surface tension govern the stability of cell shape. Our estimate for bending rigidity *κ* = 0.22 MPa *µ*m^3^ in *E. coli* cells, was notably higher than that reported for *Caulobacter crescentus* of 0.04 MPa *µ*m^3^ (*Wright, et al.* 2015). This difference could either result from differences methods of estimation, or physiological differences in cell wall and MreB mechanics. Perturbation of MreB polymerization with A22 disrupts the balance between bending rigidity counter-acting internal growth pressure, leading to the observed morphological changes.

Taken together, we conclude that bending rigidity and surface tension together integrate in cell shape and size maintenance.

Along with the mechanical shell-model, we have also invoked two phenomenological models of width dynamics-a saturation and logistic growth-both of which fit the cell width dynamics of untreated cells equally well (Figure S9). The logistic model in turn successfully captures the distinction between saturating width seen in untreated cells compared to bulging dynamics in the cell centre in simulations as a result of variation of *κ* and *γ* (Figure 6A). The estimate of a maximal width, *w_max_*, allowed us to infer a width-threshold for regulated growth (region 1), cell shape changes (region 2) and uncontrolled increase in bulge size (region 3), which in turn match well with the experimentally observed cell shapes (Figure 6D). Such a phase diagram is predictive for other bacterial cell types of rod-shape, with a boundaries changing depending on the individual mechanics of wall, membrane and cytoskeleton.

Taken together we have identified key mechanical parameters such as turgor pressure, surface tension, and bending rigidity, that regulate cell shape and size homeostasis in *E. coli*. We apply an existing cell mechanics framework that links these parameters, providing a quantitative understanding of how bacterial cells control their dimensions, and compare model predictions to cell shapes under perturbation. We identify a threshold relation between bending rigidity and surface tension required to maintain *E. coli* cell shape that can be tested in future for generality.

## Materials and Methods

### Bacterial culture and growth in presence of inhibitors

*E. coli* MG1655 were picked from single colonies growing on plates containing lysogeny broth (LB) with 1.5% agar agar (Hi-Media, Mumbai, India) and inoculated in 5 ml liquid LB. Overnight cultures grown at 37*°*C with shaking at 180 rpm (MIR-154-PE, Panasonic Corporation, Japan) were used to inoculate a 1:100 secondary culture of 5 ml of LB and grown to mid-log phase for approximately 1.5 hours. A cell suspension of ≈1 ml was sampled from the mid log-phase culture, centrifuged at 8,000 rpm for 12 minutes and the pellet resuspended in 1 ml sterile phosphate buffered saline (PBS). This was repeated three times and 10 *µ*l of the suspension placed on a coverslip, and sandwiched with 1.5% agarose pad containing LB a combination of the following inhibitors: 2 *µ*g/ml A22 (Sigma-Aldrich, Inc.) and 10 *µ*g/ml Cephalexin (Sigma-Aldrich, Inc.) in a custom-build 3D printed assembly.

### 3D printing and device assembly for live cell imaging

The structure was designed using the free web-based software Tinkercad (Autodesk, Inc., CA, USA https://www.tinkercad.com/). The assembly consisted of 2 parts: mold and holder. The design was visualized for the print view and exported to .gcode format using the program Cura ver. 2.25.0 (Ultimaker B.V., Geldermalsen, The Netherlands) and printed using a 3D printer Ultimaker 2+ (Ultimaker B.V., Geldermalsen, The Netherlands) with a 4 mm poly lactic acid (PLA) filament with printer settings of infill 60%, adhesion ON, layer height = 0.2 mm and other parameters based on the instrument defaults.

Once printed, the mold unit was placed on a 25 x 75 mm glass slide and 1 ml molten LB + 1.5% (w/v) agarose was pipetted and allowed to solidify at room temperature (Figure S1I). The mold + agarose pad assembly was inverted, covered with a 22×22 mm coverslip, and a 22×40 mm coverslip (Hi-Media, Mumbai, India) added to the base (Figure S1II). The mould assembly with two coverslips was assembled with the holder and fastener clips for stability and used further for time-lapse microscopy (Figure S1III-V).

### Microscopy and image analysis

Cells of *E. coli* MG1655 were imaged in 100x (N.A. 0.95) DIC mode using an Nikon TiE Inverted Microscope with control software NIS Elements ver.4.30.01 (Nikon Corp., Japan). Images were acquired on a CCD camera Andor Clara 2 (Andor, NI, U.K). Images were pre-processed and analyzed using a script written in MATLAB R2022a (Mathworks Inc., Natick, MA, USA) or interactively depending on the type of data, to measure cell lengths and widths with the DIC cell images as inputs. Cell outlines were segmented by detecting edges using the Canny method (*Canny*, 1986) with threshold 0.2. Gaps were filled and regions bridged to arrive at cell outlines. Cells were filtered based on a minimum solidity criterion between 0.3 and 1 to remove small and irregular objects, with cell boundaries optimized using an edge-based active contour method (*Kass, et al.* 1988), with a smoothing factor 1 and a contraction bias of −0.1. Cells with contours were oriented vertically along their major axis and the midline calculated by iteratively finding the center of mass along the vertical axis. Cell widths were taken to be the means of the 3rd and 4th quartiles of the minimal distance between the midlines and the contour boundaries. By taking only top 50% values we exclude narrow widths at the ends of the cell from analysis.

### Data analysis and fitting

#### Parsing mother-machine data

Length and width time-series of *E. coli* strains grown in the mother-machine were obtained a publicly available repository (Jun lab website) based on their previous results (*Wang, et al.* 2010). The cell division annotation (1: division, 0: cell growth) was used to obtain birth-to-division time-series for model fitting. Only those time-series with doubling time (*t_d_*) ranging between 10 ≥ *t_d_* ≥ 35 minutes, based on the distribution doubling times (*Osella, et al.* 2014), with the bounds including most of the data. We observed that there are spurious doubling times as low as 1 to 5 minutes, non-physiological (*Osella, et al.* 2014) we set the lower bound to be 10 minutes. The upper bound is decided such the most of the data is accounted for. A detailed description of data parsing is provided in the supplementary material (Figure S3).

#### Virtual synchronization of cell shape dynamics

Cells treated with a combination of A22 and Cephalexin were recorded and analyzed for cell shape. However, since the data came from multiple fields of view and varied initial time points, the bulge dynamics needed to be aligned in time. The time-series data of width was parsed in a custom written Python code, the second derivative with respect to time calculated at each time point, its maximum was used to identify the time at which width inflection occurred, *t_bulge_* = 0. This was used to align the time-series from multiple ROIs, which could then be averaged and used further to fit the ODE model.

#### Fitting phenomenological models to width dynamics

The logistic growth equation used to fit the simulated width dynamics for varying *κ* and *γ* values:

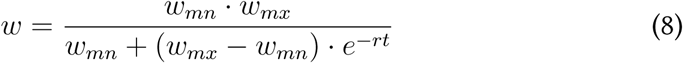

where *w_mn_* is the minimum width, *w_mx_* is the maximum width, *r* is the growth rate of width and *t* is the time. This logistic growth equation and the phenomenological model (Equation 5) was fit to the cell width dynamics data of untreated cells using custom built Python scripts (Python3) using Pandas, NumPy (*Harris* 2020), SciPy and Matplotlib (v3.4.2) (*Hunter* 2007) packages. The *polyfit* function from SciPy was used for curve-fitting which uses dog-box method to perform fitting.

### Parameter estimation from ODE model fitting by particle swarm optimization

The coupled ODE model (Equations (1) and (2)) was solved using a stiff solver *ode15s* in MATLAB R2022b (Mathworks Inc., Natick, MA, USA) with initial conditions based on average birth-length and radius from observations.

Particle swarm optimization (PSO) (*Kennedy and Eberhart*, 1995) was used to find the parameter values by minimizing the error between the binned-averaged data points and the simulation using L2 norm, using the implemented methods in the MATLAB Global Optimization Toolbox. The parameters *γ*, *κ*, *η_L_*, *η_r_*, *P* and *R*_0_ were varied within specified ranges, to find the optimum. The data of mother-machine grown *E. coli* (*Wang, et al.* 2005) were fit with lower (lb) and upper bounds (ub) of each parameter of the ODE model (Table S2. Experiments where cells were inhibited by both A22 and cephalexin, i.e. resulting in the central bugle, were quantified in terms of the width of the bulging region and the pole-to-pole length and fit using bounds different from untreated cells (Table S2, since the cellular dynamics are expected to have been altered (Fig. S8). Cells treated with inhibitors and grown on agarose pads were fit to the coupled ODE model as follows: (i) the best-fit was found for every individual cell in each treatment or shape: untreated, +Ceph, +A22, Pre-bulge, Post-bulge, Rugby, Scaled Rod, 2-Bulge,

(ii) the mean P and R_0_ values were fixed based on the ensemble average for each condition,

(iii) each cell were once again fit using these average constraints of P and R_0_ to obtain the only two free parameters - bending rigidity and surface tension (Table S2).

We required ~160 min to fit the data from ≈ 10^3^ cells using code run on a MacBook Pro with a 2.9 GHz 6-core intel i9 processor and 32 GB RAM.

## Code availability

The codes used for the different analysis methods (a) image analysis, (b) data parsing and (c) ODE model simulations and fitting are available as OpenSource. These are deposited in a GitHub repository https://github.com/CyCelsLab/bactShapedyn/.

## Author Contributions

Tanvi Kale carried out all experiments, image analysis and analyzed the data. Ryth Dasgupta carried out the model simulations, fitting and analyzed the data. Mandar M. Inamdar advised us on the model solution and simulations. Chaitanya A. Athale designed the study and supervised the work. Tanvi Kale, Ryth Dasgupta and Chaitanya A. Athale wrote the article.

## Declaration

The authors declare no competing interests.

## Acknowledgments

TK is funded by a PhD fellowship from IISER Pune. RD is supported by a fellowship from the Department of Science and Technology, Govt of India. This work is supported by Grant BT/PR40262/BTIS/ 137/38/2022 from the Dept. of Biotechnology, Govt of India to CAA. We are grateful to Aman Soni for data-analysis inputs, Sunish Radhakrishnan for access to a phase-contrast microscope and Nishad Matange for the kind gift of bacterial strains.

## Supporting Material

- Supplementary Tables
- Supplementary Figures
- Supplementary Movies
- Supplementary Methods

## Supplementary Tables

**Table S1:**
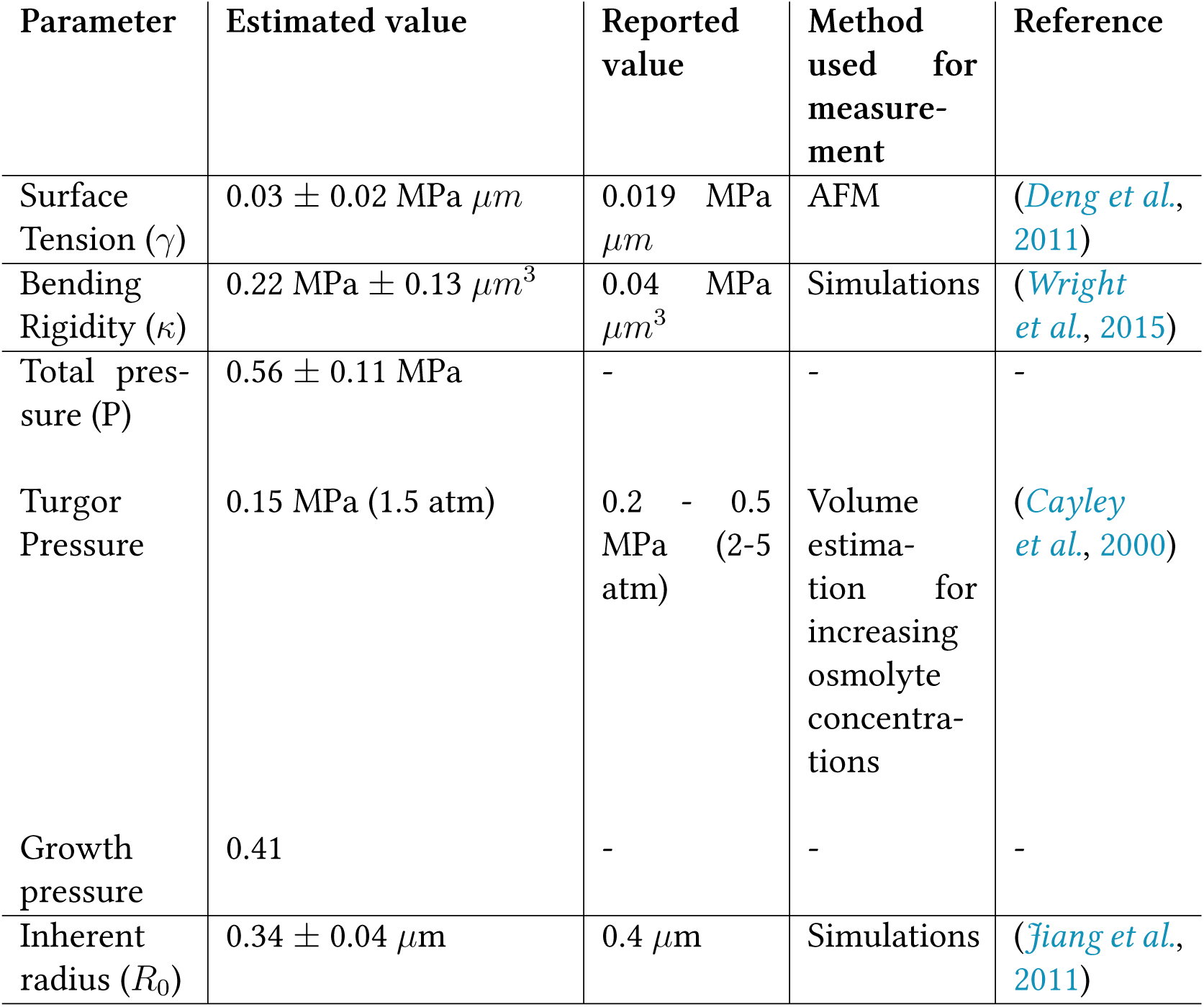
Comparing estimated mechanical parameters to previous reports. Estimates of the surface tension (*γ*), bending rigidity (*κ*), pressure (P) and inherent radius (R_0_) obtained from the ODE model are compared to previous reports from literature. Each value is the mean ±s.d. obtained by averaging the mean values from fits to *E. coli* strains SJ108, SJ119, CGSC 6300 in Figure 2D.

**Table S2:** Criteria provided to the optimization routine to fit the coupled ODE model to multiple experimental datasets. The upper and lower bounds provided to the PSO algorithm in order estimate the mechanical parameters *γ*: surface tension; *κ*: bending rigidity of the envelope; *η_L_* and *η_w_*: viscosity in length and width respectively; P: total pressure; *R*_0_: equilibrium radius of the spherocylinder.

| Data | Iteration | Upper (UB)/<br>Lower (LB)<br>bound | $\gamma$<br>(MPa<br>$\mu m$) | $\kappa$<br>(MPa<br>$\mu m^3$) | $\eta_L$<br>(MPa<br>s) | $\eta_w$<br>(MPa<br>s) | P<br>(MPa) | $R_0$<br>($\mu m$) |
| --- | --- | --- | --- | --- | --- | --- | --- | --- |
| Jun Lab | I | LB | 0.01 | 0.12 | 500 | 2000 | 0.3 | 0.1 |
|  |  | UB | 0.12 | 0.8 | 1800 | 4000 | 0.7 | 0.5 |
|  | II | LB | 0.01 | 0.12 | 700 | 2500 | 0.3 | 0.1 |
|  |  | UB | 0.12 | 0.8 | 700 | 2500 | 0.7 | 0.5 |
| Perturbations | I | LB | 0.01 | 0.01 | 1200 | 1000 | 0.2 | 0.1 |
|  |  | UB | 0.08 | 0.2 | 1800 | 4000 | 0.7 | 0.5 |
| | II | LB | 0.01 | 0.12 | 1500 | 4000 | $P_{avg}$ | $R_{0avg}$ |
| | | UB | 0.12 | 0.8 | 1500 | 4000 | $P_{avg}$ | $R_{0avg}$ |
| Post bulge | - | LB | 0.01 | 0.12 | 500 | 500 | 0.2 | 0.1 |
|  |  | UB | 0.12 | 0.8 | 8000 | 4000 | 0.7 | 0.5 |

**Table S3:** Comparing the parameters obtained by fitting width growth dynamics to the Saturation width model (model III) (Equation 5)and the logistic growth equation (Equation 8) The parameters obtained from the saturation width model are *w_mn_*: minimum width, *w_mn_*: maximal width and *t*_1_*_/_*_2_: time at which the width is half maximal are compared to the parameters obtained from the logistic growth equation, where *w_mn_*: minimum width, *w_mn_*: maximal width and *r*: rate of width growth.

| Parameters | Description | Saturation width model (model III) | Logistic growth equation |
| --- | --- | --- | --- |
| $w_{mn}$ | Minimum width | 0.62 $\mu\text{m}$ | 0.63 $\mu\text{m}$ |
| $w_{mx}$ | Maximum width | 0.75 $\mu\text{m}$ | 0.74 $\mu\text{m}$ |
| $t_{1/2}$ | Time at which width is half its maximum value | 3.75 min | - |
| $r$ | Rate of width growth | - | 0.2 $\mu\text{m}/\text{min}$ |

## Supplementary Figures

**Figure S1:**
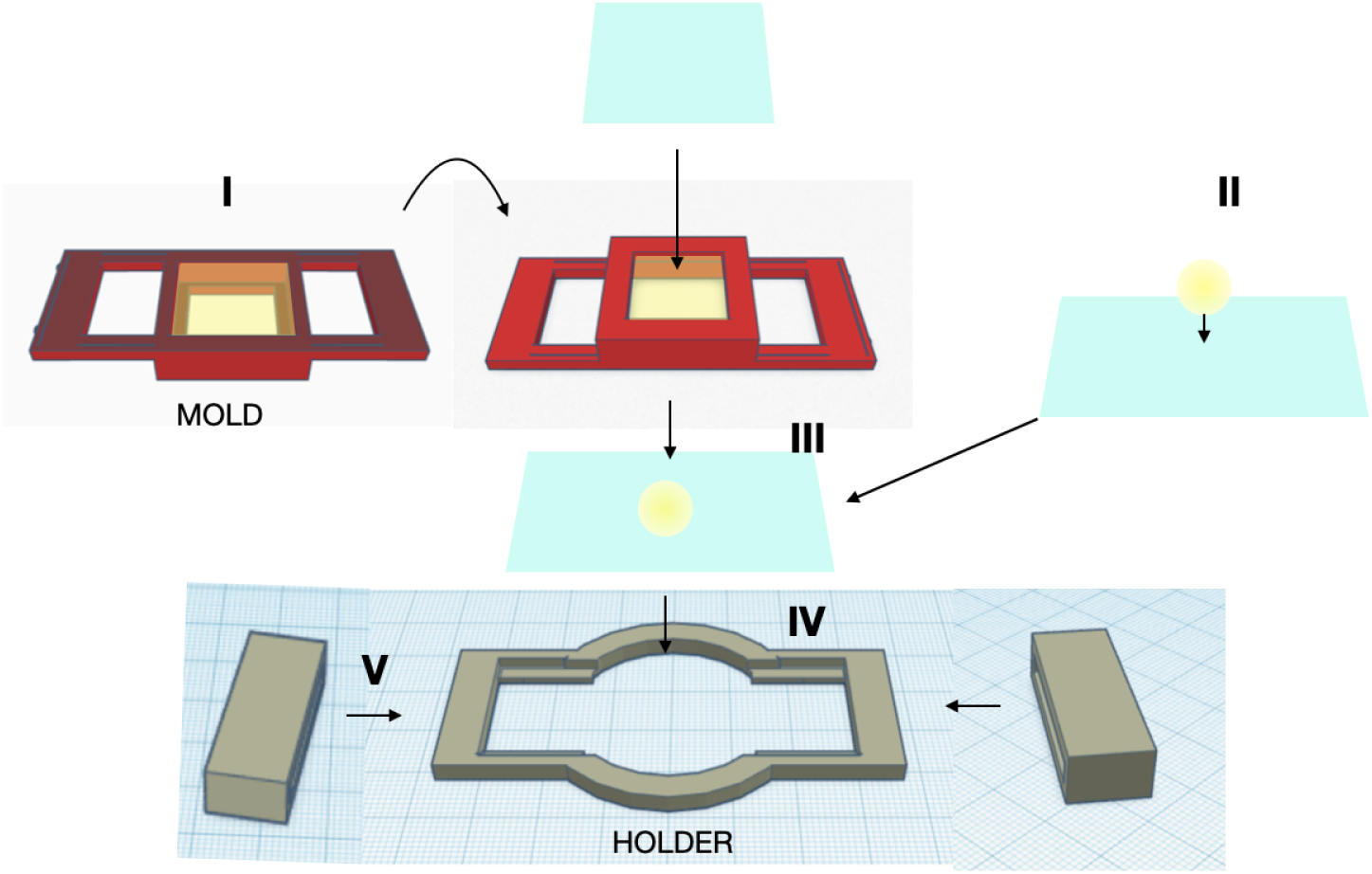
Schematic representation of 3D printed holder for live cell imaging. I: The 3D printed mold was placed on a glass side and 1 ml molten LB-agarose (1.5% w/v) pipetted into it. II. 10 *µ*l culture from mid-log phase was added to a 40 x 22 mm coverslip (c.s.). III. The mold was assembled with the c.s. with bacteria to combine the setup. IV: The mold+c.s. was then placed on the 3D printed holder and V: 3D printed clips were slid on to the combined assembly, to prevent movement of components when mounted on the microscope. Arrows: sequence of assembly.

**Figure S2:**
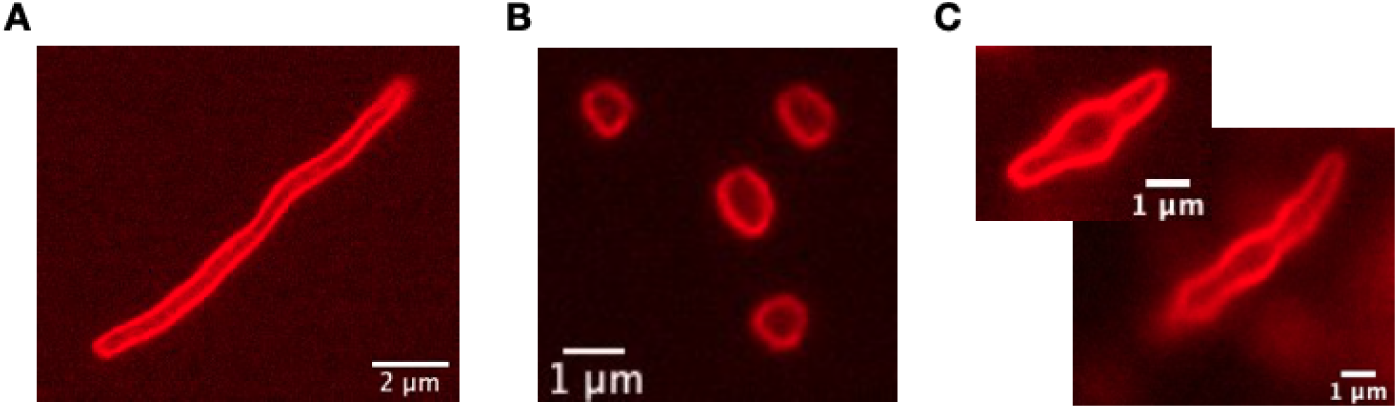
Membrane label of cells inhibited by A22 and cephalexin. Cells treated (A) 10 *µ*g/ml cephalexin (B) 2 *µ*g/ml A22 or (C) a combination of both labelled with 2 *µ*g/ml FM4-64 to visualize the membrane in fluorescence using epifluorescence microscopy (TRITC channel). Scale bar: 1 *µ*m.

**Figure S3:**
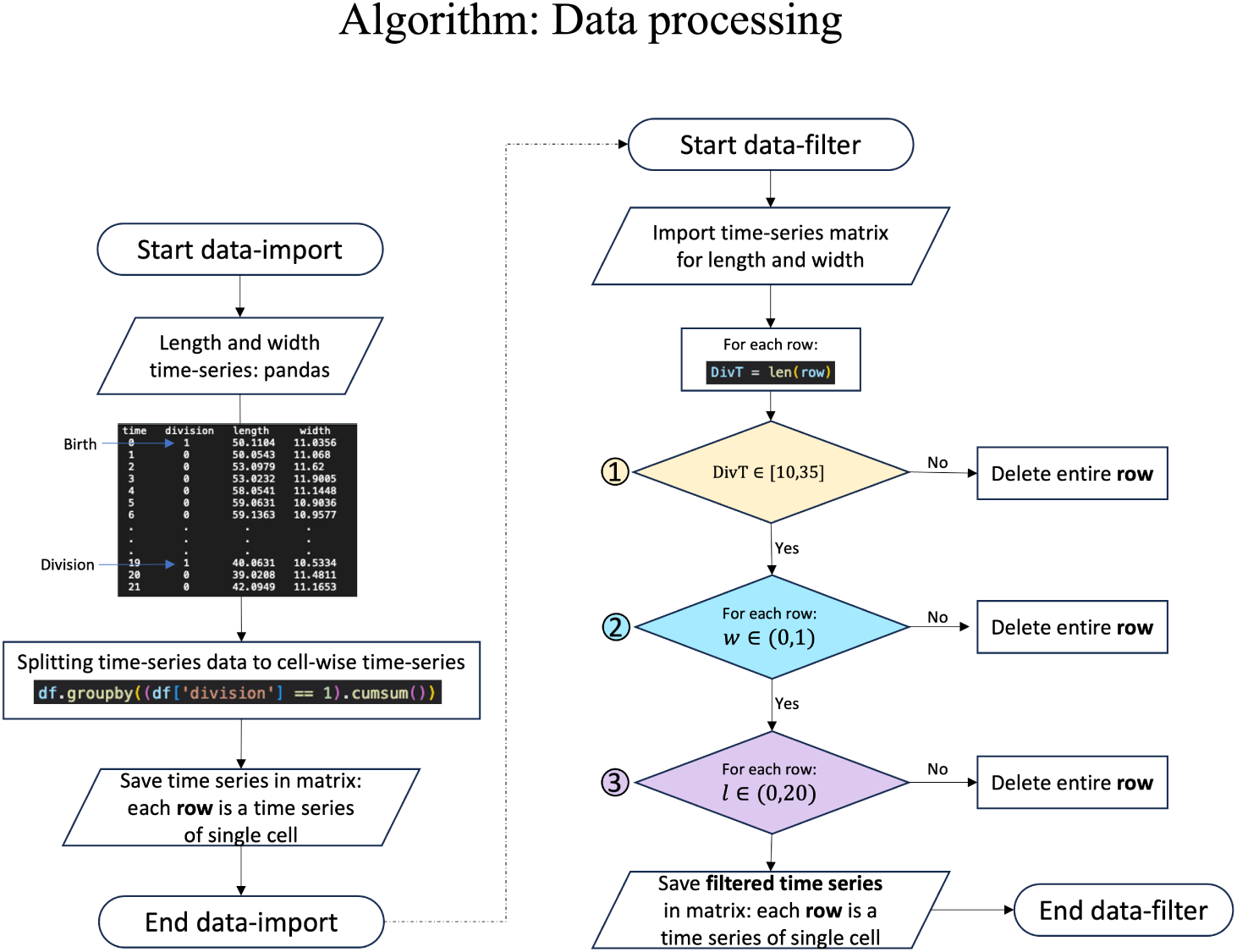
Schematic representation of the steps involved in parsing and pre-processing the data of length and width dynamics from previous work (*Wang, et al.* 2005).

**Figure S4:**
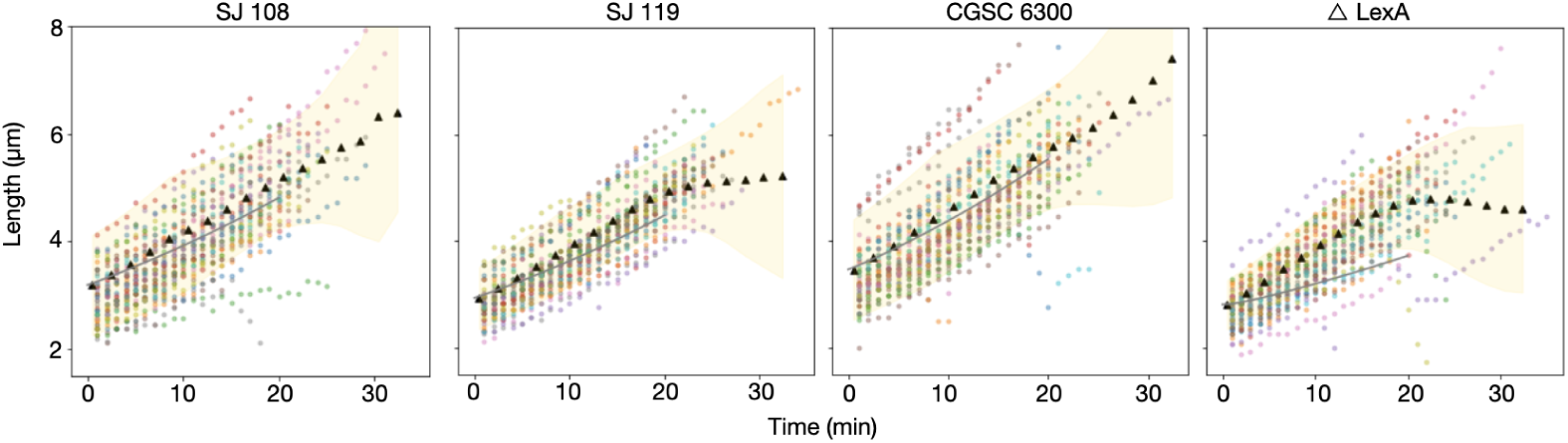
Mechanical model fit to length dynamics. The length dynamics data from (*Wang, et al.* 2010) was used to fit the coupled ODE model of length and width dynamics (Figure 2).

**Figure S5:**
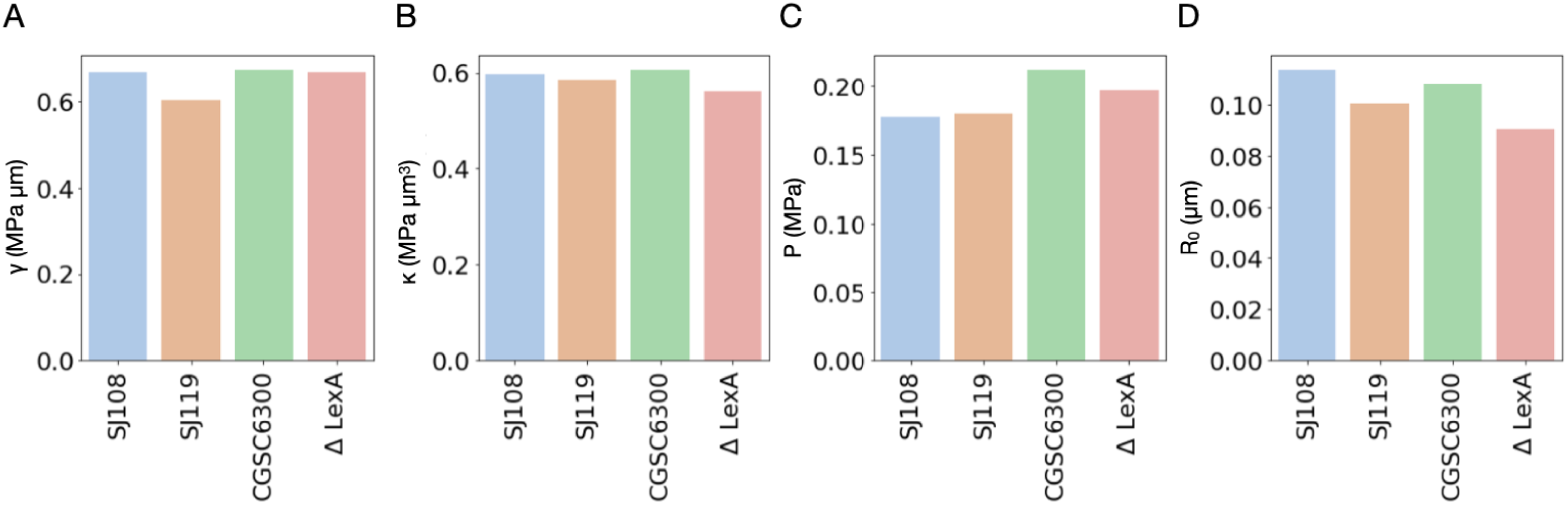
Coefficient of variation for model fit parameters for different strains. CV (Standard deviation/mean) calculated of the estimated parameters from model fitting are compared across strains (colors) in terms of surface tension, (A) surface tension, *γ* (MPa *µ*m); (B) bending rigidity, *κ* (MPa *µm*^3^); (C) total pressure, P (MPa) and (D) equilibrium cell curvature, *R*_0_ (*µ*m).

**Figure S6:**
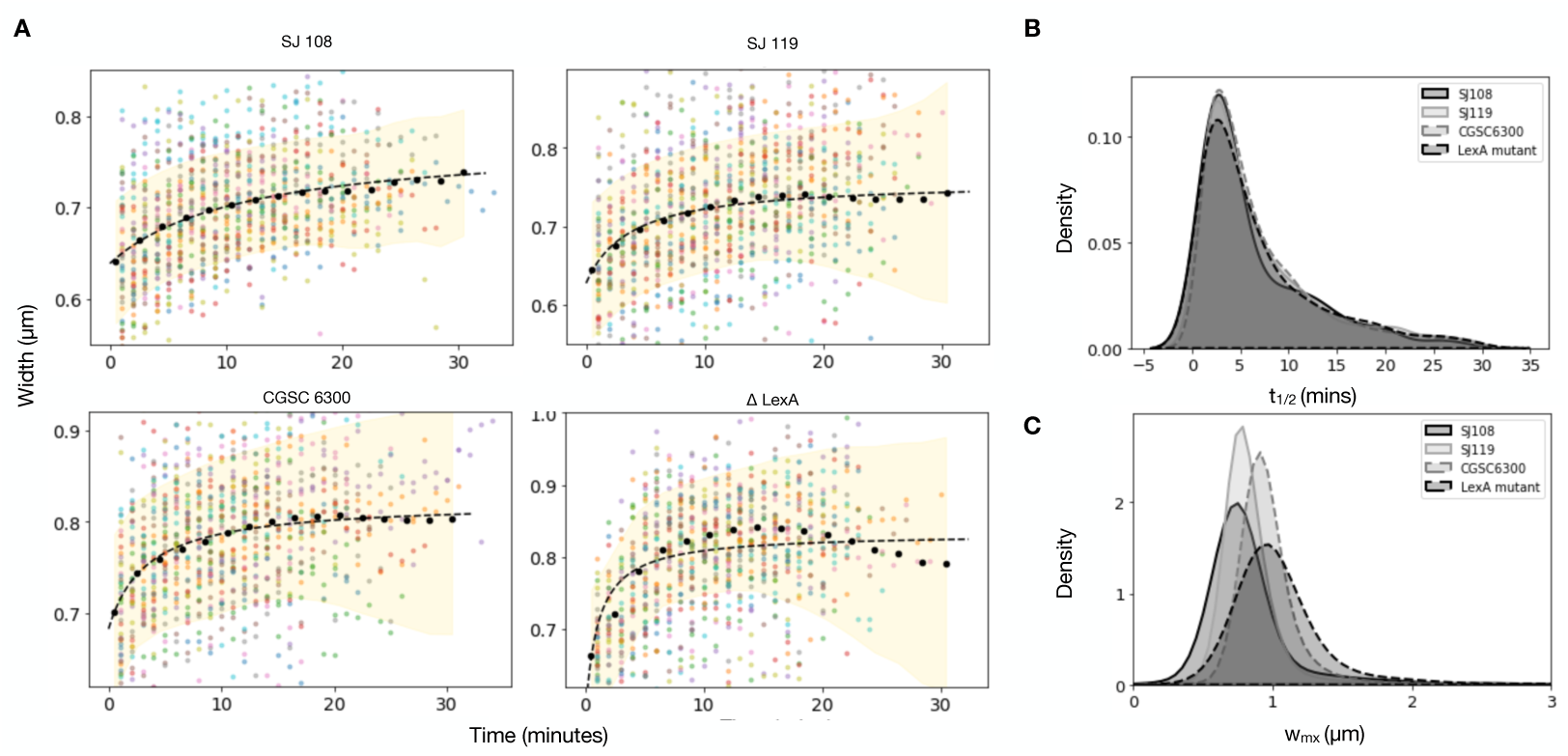
**Width saturation dynamics model fit to** *E. coli* **microfluidics data to estimate half-time to saturation**, *t*_1/2_ **and maximal width**, *w_mx_*. (A) Cell width dynamics with time from individual cells are plotted (circles) and binned average dynamics (•) ± s.d. (shaded area) were fit the saturation model (III, Equation (5)) for multiple *E. coli* strains SJ108, SJ119, CGSC6300, ΔLexA, using previously published data (*Wang, et al.* 2010). (B) The half-time to saturation, *t*_1_*_/_*_2_, and (C) maximal width, *w_mx_*, value distributions are obtained fits to individual cells. The density is plotted using a kernel distribution estimation (KDE). The shades indicate *E. coli* strains.

**Figure S7:**
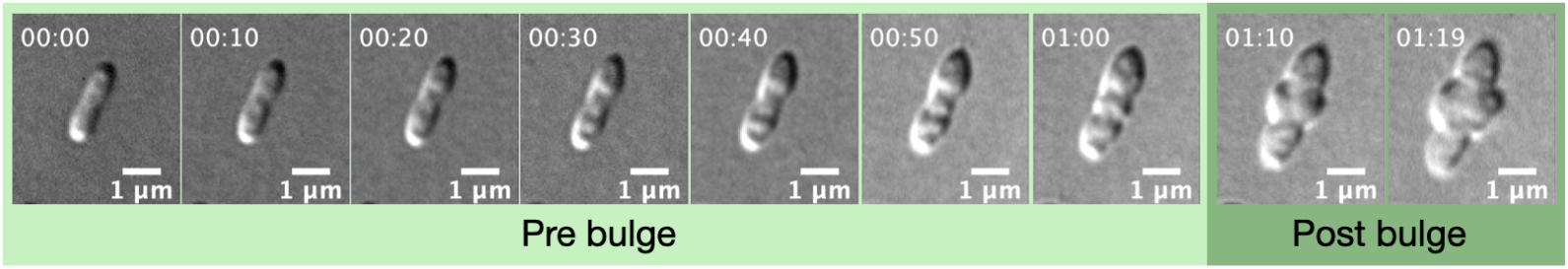
Montage of a growing cell treated with a combination of A22 and cephalexin. *E. coli* treated with 2 *µ*g/ml A22 and 10 *µ*g/ml Cephalexin over 120 minutes. Colors represent different conditions: Pre bulge (light green) and Post bulge (dark green).

**Figure S8:**
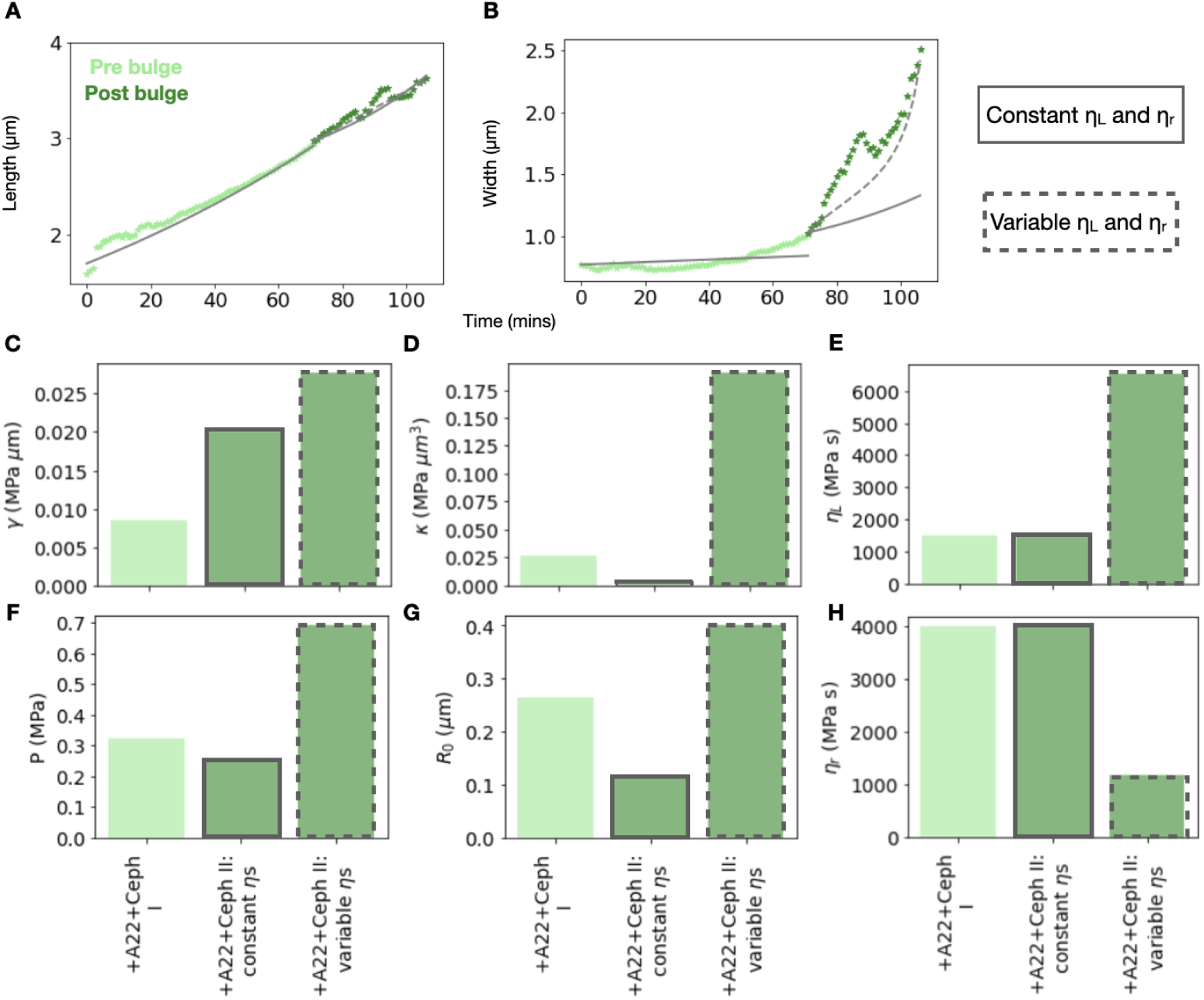
Effect of changing effective envelope viscosity in length (*η_L_*) and width (*η_w_*) (A) The cell length and (B) width dynamics of cells as a function of time from *E. coli* MG1655 treated with a combination of A22 and cephalexin (+) were fit by the mechanical model of cell growth either with fixed values of *η_L_* and *η_w_* (gray lines) or these values were optimized (dashed-line). (B) (C-H) Bar charts showing values obtained from PSO for different conditions for (C) Surface tension, *γ* (MPa *µ*m) and (D) Bending rigidity, *κ* (MPa *µm*^3^), (E) Viscosity in Length, *η_L_* (MPa s), (F) Total pressure, P (MPa), (G) equilibrium cell curvature, *R*_0_ (*µ*m) and (H) Viscosity in width, *η_w_* (MPa s). Colors represent different conditions: +A22+Ceph I - Pre bulge (light green), +A22+Ceph II - Post bulge (dark green) which is divided further into two parts: constant *η_L_* and *η_w_* (solid line) and variable *η_L_* and *η_w_* (dashed line).

**Figure S9:**
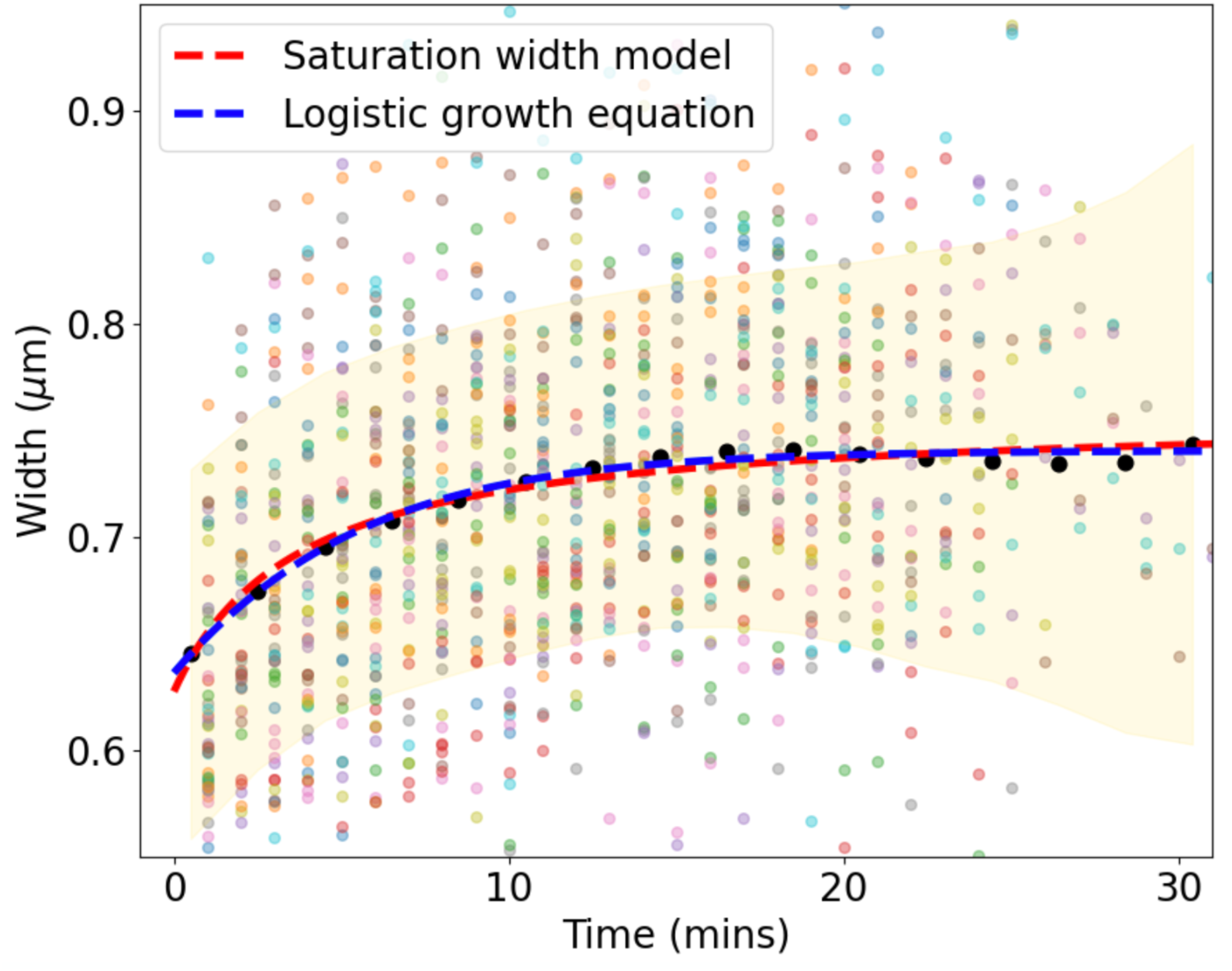
Comparing the fits for the saturation width model and the logistic growth equation to width dynamics data. Cell width dynamics with time from individual cells are plotted (circles) and binned average dynamics (•) ±s.d. (shaded area) were fit the logistic growth equation (Equation (8)) and the saturation width model (III, Equation (5)) for *E. coli* SJ119 from previously published data (*Wang, et al.* 2010) using the parameters described in Table S3.

## Supplementary Videos

**Video SV1:**
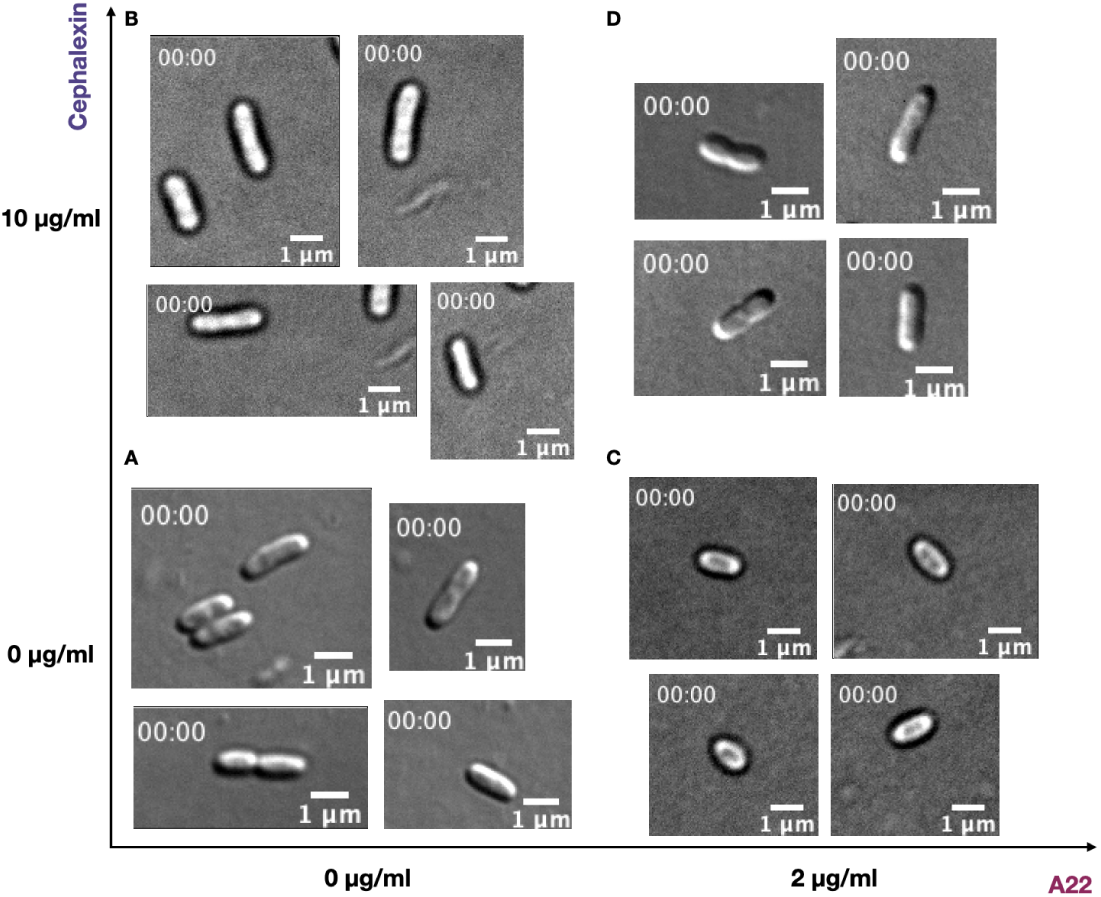
E. coli perturbed with A22 and cephalexin. Representative ROIs of DIC time series (100x) of *E. coli* (A) MG1655 untreated (bottom left), (B) treated with 10 *µ*g/ml cephalexin (top left), (C) 2 *µ*g/ml A22 (bottom right) and (D) a combination of both (top right) Scalebar: 1*µ*m, Time interval: 1 minute between 2 frames.

## Supplementary Methods

### Parsing single-cell growth data obtained from published microfluidics experiments

Publicly available data from the Jun lab website (*Wang, et al.* 2005) was used to fit the model. We followed a simple algorithm to import and filter the data to remove outliers based on physiological constraints. The relevant information from the downloaded data were the ‘Time’, ‘Length’, ‘Width’ and ‘Division’ columns.

Four different strains of asynchronous cell cultures were measured using a mother machine. The measurement of a strain had data imaged (and then analyzed) from multiple regions of interest (ROI). Each ROI had multiple channels for the cells to grow in, and each channel had three to four cells.

We explain the steps of the algorithm in a little more detail in the following subsection:

### Steps used to filter the data

1. Import raw data into python using pandas library
2. Save the data in a pandas data frame
3. Split the time-series data into cell-wise time-series data. This is done by treating all data values sandwiched between two division events (marked by the division boolean = 1) as the chronological data values for a single cell.
4. These cell-wise time-series data are saved in rows of a matrix. The length of these rows signifies how long the cells have grown for before dividing.
5. The cell-wise data is taken to the data-filter algorithm which removes entire cells that have values outside the allowed limits.
6. The doubling time for each cell is calculated by the length of data-series for that cell.
7. All cells that had doubling times less than 10 or more than 35 minutes (both exclusive) were discarded. These limits were chosen after inspecting the distribution of the doubling times. (a) We observed that there are spurious doubling times as low as 1 to 5 minutes, non-physiological (*Osella, et al.* 2014) we set the lower bound to be 10 minutes. The upper bound is decided such the most of the data is accounted for.
8. Since the channel size of the mother machine used to make the time-series measurements was 1 *µ*m, it could not have been possible for cell widths to be more than 1*µm*. So, all cells that had widths that were greater than or equal to 1 *µ*m were discarded. Also, the width could not have been zero, so cells that had 0 width were also discarded.
9. A similar thing was done for lengths. Cells that had lengths less than or equal to 0, and greater than or equal to 20 *µ*m were discarded. The reason we chose 20 *µ*m as the upper bound, was because the cells appeared filamentous, and so we did not have a physiological upper bound for length. We went with 20 *µ*m arbitrarily, since it included most of the cells.
10. Finally the filtered cell time-series are exported for further analysis.

Having cleaned the data, we binned the data into 17 bins, each with a width of 2 minutes. All the data points for width and length between 0 and 2 minutes were assigned to bin 1, points between 2 and 4 minutes were assigned to bin 2 and so on. These points were used for optimization and fitting to reduce the computation time since dealing with fewer points would be faster than dealing with all the points.

## Notes

### Competing Interest Statement

The authors have declared no competing interest.

### Summary of Updates

Some text has been edited for correctness.

